# Modern human expansion from (southern?) Africa: Origin, non-African ancestry, and linearity

**DOI:** 10.1101/2022.07.31.500977

**Authors:** Zarus Cenac

**Affiliations:** Department of Psychology, City, University of London, Rhind Building, St John Street, London, United Kingdom, EC1R 0JD

## Abstract

Previous research favours the idea that modern humans spread worldwide from Africa. For instance, through autosomal diversity, a geographical area of origin for this worldwide expansion is indicated to entirely be within Africa. It remained to be seen if this indication happens for certain variables such as Y-chromosomal diversity. There is disagreement regarding where in Africa the origin is. Perhaps a region of Africa may seem to be the origin because of non-African ancestry rather than the expansion. The present research considered whether some genetic and cranial variables indicate the expansion, and, furthermore, where in Africa the expansion started. Variables included, for example, autosomal diversity (in sub-Saharan Africa) which was adjusted for non-African ancestry, Y-chromosomal diversity, and cranial sexual size dimorphism. For each variable, it was seen if the estimated area of origin was solely in Africa. Moreover, to generally estimate the origin, a centroid was calculated from potential origins which were obtained in the present, and past, research. The area of origin was completely within Africa for each variable except one – for Y-chromosomal diversity, the area was possibly in Asia only. The centroid of the potential origins was in southern Africa, consequently supporting a southern African origin. The autosomal diversity of sub-Saharan African populations, adjusted for non-African ancestry, indicated a southern African origin. The adjusted diversity appeared to start declining from about 2,000-3,000 km away from the origin – this non-linear pattern may be explained by travelling which happened after the expansion, or the expansion entering locations in Africa which were already populated, to differing extents, by modern humans.

## Introduction

Findings in the literature align with modern humans having journeyed across the world via an expansion that originated in Africa (Manica et al., 2007; Ramachandran et al., 2005; see Henn, Cavalli-Sforza, & Feldman, 2012, for a review). Research agrees with population bottlenecks having been encountered during the expansion (more and more bottlenecks with greater and greater distance from Africa) thereby leading to a reduction in autosomal microsatellite diversity (expected heterozygosity) as the expansion reached greater distances (Prugnolle, Manica, & Balloux, 2005). Not only does the diversity of autosomal microsatellites decrease as geographical distance from Africa widens (Prugnolle, Manica, & Balloux, 2005), but declines are also present in other aspects of biological diversity, such as diversity in autosomal single nucleotide polymorphism (SNP) haplotypes (Balloux et al., 2009) and cranial shape (von Cramon-Taubadel & Lycett, 2008).^1^

Through measuring the fit between geographical distance (the distance from geographical locations to populations) and a biological variable (e.g., the autosomal microsatellite diversity of populations), it is possible to estimate a geographical area in which the expansion started (Manica et al., 2007). This area contains the location which gives the best fit, and locations which are similar in fit to the best (e.g., Manica et al., 2007) – the area can be called a **geographical area of best fit** (see Figure 1A for an example, and the *Glossary* in Box 1). More than one geographical area of best fit may be found with a biological variable (as in the present research with mitochondrial diversity, and also with cranial shape diversity for female crania). For instance, one geographical area of best fit may have correlation coefficients (for relationships between distance and diversity) that are negative, whereas another geographical area of best fit has positive coefficients (see *Results and discussion*).^2^ Results in prior research have been in alignment with types of biological diversity declining as distance increases from the origin of the expansion (e.g., Ramachandran et al., 2005; von Cramon-Taubadel & Lycett, 2008), so the geographical area of best fit which has negative correlation coefficients (rather than the one with positive coefficients) would be favoured for being the area where the expansion has its origin.

**Figure 1.**
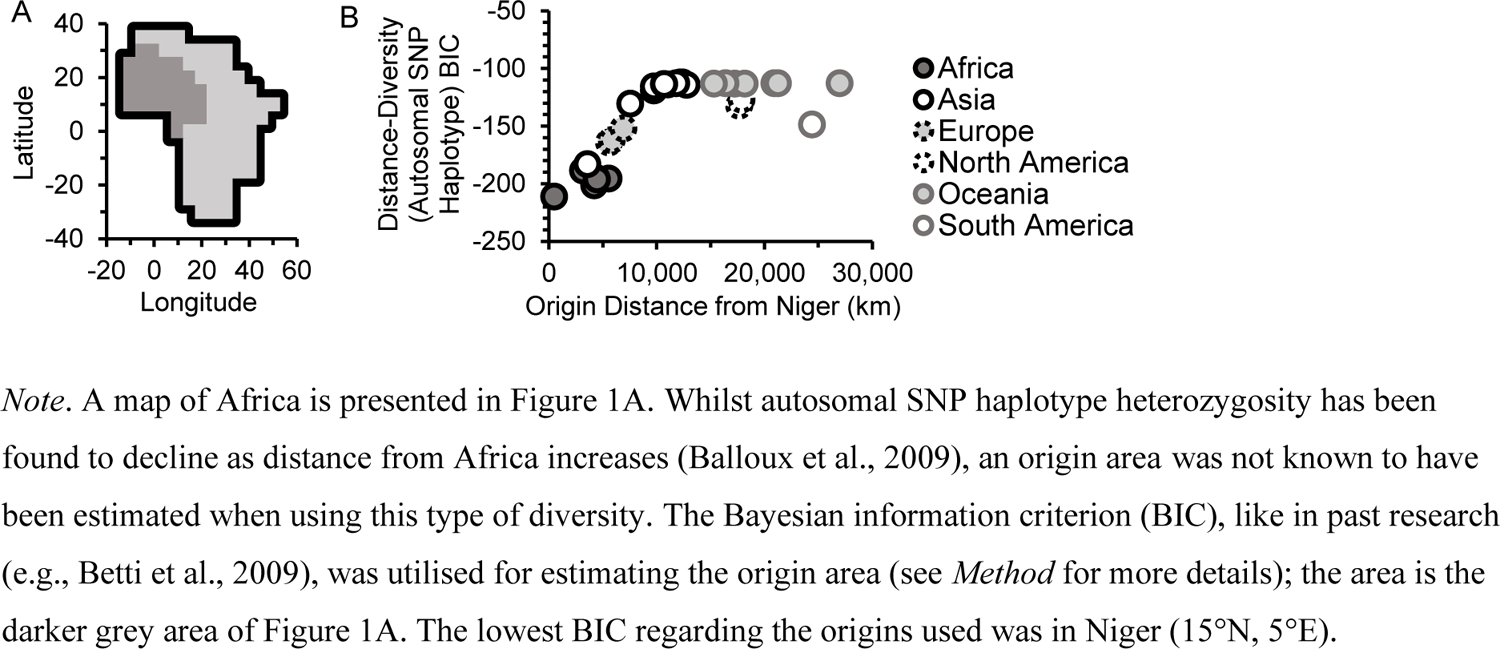
Autosomal SNP Haplotype Diversity Suggests the Worldwide Expansion Originated in Africa

### Skeletal and genetic

A variable which appears to mark the expansion from Africa i) relates to distance from Africa (e.g., Prugnolle, Manica, & Balloux, 2005; von Cramon-Taubadel & Lycett, 2008). Moreover, like autosomal microsatellite diversity (Manica et al., 2007), a variable which indicates the expansion should ii) lead to an estimated **origin area** entirely within Africa. Ideally, an indicator of the expansion should meet both of these points (i.e., association with distance from Africa, and the estimated area being in Africa alone).^3^ Of the three studies on skeletal diversity which have indicated origin areas (Betti et al., 2009, 2013; Manica et al., 2007), only cranial form diversity amongst males (e.g., using 105 populations [*n* = 4,666]) has fulfilled both points (Betti et al., 2009; Manica et al., 2007) – the area is fully within Africa when adjusting the cranial diversity of males for climate (Manica et al., 2007) and without amending for climate (Betti et al., 2009) (see Manica et al., 2007, and Betti et al., 2009, for methodological differences between the two studies). Female cranial form diversity has met point one but not point two (Betti et al., 2009), and the same can be said for the hip bone shape diversity of males and females (Betti et al., 2013).^4^ Other studies on whether distance from Africa is related to types of skeletal diversity (e.g., Betti et al., 2012), such as cranial shape diversity (von Cramon-Taubadel & Lycett, 2008), have not found origin areas.^5^

In unpublished research, cranial sexual size dimorphism (adjusted for absolute latitude) seemed in line with being an indicator of expansion from Africa (Cenac, 2022). The adjusted dimorphism was found to attain point one, i.e., it correlated with distance from Africa (and correlation coefficients looked like they were at their numerical highest when employing distance from Africa) (Cenac, 2022). Nonetheless, research was not known to have tested if it meets the second point, and therefore finding the origin area through this adjusted dimorphism is something which could be explored (Cenac, 2022).

As geographical distance from Africa grows, the diversity (heterozygosity) of autosomal microsatellites decreases linearly (Prugnolle, Manica, & Balloux, 2005). Using autosomal microsatellite heterozygosity, the **area of origin** is estimated to solely be within Africa (Manica et al., 2007). Aside from autosomal diversity, does any other type of genetic diversity i) decline with rising distance from Africa (Balloux et al., 2009; Shi et al., 2010) and ii) result in Africa completely having the estimated origin area?

Balloux et al. (2009) found declines (as distance from Africa increases) in X-chromosomal microsatellite heterozygosity, Y-chromosomal microsatellite heterozygosity, and mitochondrial diversity.^6^ Figure 3E In Balloux et al. (2009) presents Y-chromosomal microsatellite diversities for populations at distances from Africa. That figure (in Balloux et al.) may indicate that Y-chromosomal microsatellite diversity is not at its highest in Africa. Indeed, when diversities are grouped by continent and averaged, the numerically highest diversity is not found to be with Africa, but with Asia (Table 1). When research on diversity or/and expansion (Manica et al., 2007; Ramachandran et al., 2005; Romero et al., 2009) is considered, one would expect Y-chromosomal diversity to numerically be greater in Africa than anywhere else if Y-chromosomal diversity is a reflection of the expansion from Africa. Therefore, Table 1 seems at odds with Y-chromosomal diversity actually indicating the *worldwide expansion*.^7^ Given Table 1, it would not be surprising if Y-chromosomal microsatellite diversity leads to an origin area which is outside of Africa. Elsewhere, in Chiaroni et al. (2009), there was a negative gradient regarding the Y-chromosomal haplogroup diversity of populations and distance from Africa. Yet Figure S6 of that study (i.e., Chiaroni et al., 2009) may suggest that this diversity is not generally at its greatest in Africa.

**Table 1.** Average Y-Chromosomal Microsatellite Diversity.

| Continent | Y-Chromosomal Diversity |
| --- | --- |
| Africa | 0.42 |
| Asia | 0.46 |
| Europe | 0.44 |
| North America | 0.37 |
| Oceania | 0.39 |
| South America | 0.19 |

*Note.* Using heterozygosities within each of 51 populations in Balloux et al. (2009), averages (means) were calculated by continent. Also, see estimates of Y-chromosomal diversity elsewhere (e.g., Hammer et al., 2001; Seielstad et al., 1999), for instance, in Jorde et al. (2000), microsatellite diversity was numerically higher in Africa than in Asia or Europe (other continents were not featured).

Balloux et al. (2009) did not explore where origin areas are located regarding genetic diversities; they did not determine if mitochondrial, and X- and Y-chromosomal diversities suggest an origin area entirely within Africa. Other studies have occurred concerning distance from Africa and the diversity of the X chromosome (Li et al., 2008) and Y chromosome (Chiaroni et al., 2009; Shi et al., 2010), regarding X-chromosomal haplotypes (Li et al., 2008), and Y-chromosomal haplogroups (Chiaroni et al., 2009; Shi et al., 2010). However, these studies also did not estimate origin areas from X- and Y-chromosomal diversities. Consequently, it was of interest to estimate origin areas through X-chromosomal, Y-chromosomal, and mitochondrial diversities.

By no means has discussion on the expansion been limited to biological diversity (e.g., Henn, Cavalli-Sforza, & Feldman, 2012). Language has featured in the discourse regarding the expansion from Africa, with reference being made to Atkinson (2011) (e.g., Atkinson, 2012; Henn, Cavalli-Sforza, & Feldman, 2012). In Atkinson (2011), it was found that phonemic diversity falls as distance from Africa rises, and that the geographical area of best fit for the negative association between distance and phonemic diversity is only within Africa. Yet, elsewhere, the geographical area of best fit was outside of Africa (Creanza et al., 2015). (For discussion regarding Atkinson, 2011, see Atkinson, 2012, or Van Tuyl & Pereltsvaig, 2012, for example.)

### Origin of expansion

Where in Africa did the expansion originate? Inferring from previous research/reasoning (Henn, Cavalli-Sforza, & Feldman, 2012; Manica et al., 2007; Ray et al., 2005; Tishkoff et al., 2009), the answer is still not clear. For instance, eastern Africa has been suggested based on simulations with respect to autosomal microsatellites (Ray et al., 2005),^8^ whereas southern Africa was favoured in a review of the literature (Henn, Cavalli-Sforza, & Feldman, 2012).^9^ Figure 1 of Batini and Jobling (2011) summarised some findings in research concerning human origins onto two maps of mainland Africa. This led to an idea, in the current analyses, of gathering coordinates at which relationships between geographical distance and indicators of the expansion from Africa are at their strongest in Africa (such coordinates are called **peak points** in the present research), and showing their locations through a representation of mainland Africa. It could be seen if peak points (for different types of indicators of the expansion) generally cluster in any particular region(s); a clustering in a region would support the expansion having its origin in that region. Moreover, a centroid could be calculated from the peak points (found with indicators) as a general approximation of where the expansion started. However, the centroid may not necessarily be useful. For example, if the peak points were not clustered in a region, but dispersed across Africa, a centroid might not be indicative of which region(s) the expansion originated in. Therefore, clustering and the centroid would both be of interest.

### Non-African ancestry

The amount of non-African ancestry in African populations is a hurdle for studying expansion from Africa (López et al., 2015). Indeed, extrapolating from previous research regarding phonemic diversity (Van Tuyl & Pereltsvaig, 2012) and admixture (Gallego Llorente et al., 2015; Haber et al., 2016; Pickrell et al., 2014; Pickrell & Reich, 2014), the non-African ancestry in Africa could affect where peak points are.^10^ Moreover, because peak points help determine the origin area (e.g., Manica et al., 2007), non-African ancestry may therefore affect the estimation of origin areas. Therefore, not accounting for non-African ancestry (e.g., Betti et al., 2009; Manica et al., 2007) could mean that origin areas and peak points lack precision in their ability to reflect the expansion. The present section next considers i) if peak points are driven by African populations and ii) if non-African ancestry in Africa has an involvement in the estimation of the origin (see below).

### African populations and peak points

The location leading to the lowest negative association between geographical distance and phonemic diversity in Atkinson (2011) is in Africa (Van Tuyl & Pereltsvaig, 2012). However, if populations are grouped by geographical region (e.g., North America, South America, etc.), it can be seen that adjusting the *origin* in Africa leads to a shift in the correlation between distance and phonemic diversity amongst populations in Africa, but no shift amongst populations in regions outside of Africa (Van Tuyl & Pereltsvaig, 2012). Indeed, the location producing the strongest negative correlation only shifts nominally when using worldwide populations compared to populations in Africa alone (Van Tuyl & Pereltsvaig, 2012); populations outside Africa are immaterial to defining which particular location in Africa has the lowest negative association between distance and phonemic diversity – only populations within Africa define the location (Van Tuyl & Pereltsvaig, 2012), or so it may appear (see *Results and discussion*). Because there is a nominal shift (Van Tuyl & Pereltsvaig, 2012), maybe it could be said that the peak point (for populations worldwide) with phonemic diversity is not entirely due to populations in Africa, but perhaps largely (cf. Van Tuyl & Pereltsvaig, 2012). It stands to reason that peak points for biological indicators of the expansion should be influenced in the same way as the peak point for phonemic diversity (see *Results and discussion*).

### Non-African ancestry in Africa, and the origin

Could it be that, prior to more contemporary times, e.g., in the Bantu expansion, there was no one large expansion in Africa like what happened elsewhere (Khan, 2018)? The pattern of heterozygosity falling as geographical distance from a population increases does occur in a scenario in which there are a small number of bottlenecks (two) and admixture (with the population not being used as the source of a worldwide expansion) (Pickrell & Reich, 2014). It can be understood, from previous research (Pickrell & Reich, 2014), that these bottlenecks in this scenario correspond to locations outside of Africa. Hence, support for any area in Africa being the origin of the expansion (over other places in Africa) (e.g., Manica et al., 2007) could occur even in the absence of bottlenecks within Africa, and instead be the result of admixture.^11^ Indeed, previous research is congruent with heterozygosity decreasing with greater Eurasian ancestry amongst Africans who are admixed (Haber et al., 2016). Therefore, it can be supposed that, more generally, the heterozygosity of Africans would decrease with increasing non-African ancestry (given the fall in autosomal heterozygosity with distance from Africa in Prugnolle, Manica, & Balloux, 2005). Moreover, if genetic diversity is linked to cranial diversity (von Cramon-Taubadel, 2019), perhaps there would be relationships between non-African ancestry and diversity beyond heterozygosity, e.g., with cranial diversity. So, non-African ancestry in Africa could plausibly affect origin areas, no matter whether those areas are found via genetic or cranial data. If peak points for biological variables are driven by African populations (*Results and discussion*; cf. Van Tuyl & Pereltsvaig, 2012), then non-African ancestry within Africa could greatly be determining peak points.

### Analyses

In the present research, something was defined as being an indicator of the expansion if it met the two points raised earlier, i.e., i) it is related to distance from Africa, and ii) it produces an origin area only in Africa. The present analyses estimated where in Africa the worldwide expansion began. It was established if origin areas are completely in Africa for diversity in cranial shape, autosomal SNP haplotypes, mitochondrial DNA, X-chromosomal microsatellites, and Y-chromosomal microsatellites. Given that these cranial/genetic diversities do have negative correlations with distance from Africa (Balloux et al., 2009; von Cramon-Taubadel & Lycett, 2008), the expansion from Africa would be signalled by these diversities if the estimated origin areas are only within Africa. Additionally, more of an insight was sought into whether cranial form diversity is indicative of the expansion from Africa. Following on from previous research (Cenac, 2022), analyses found whether cranial size dimorphism (whether adjusted for absolute latitude or not) indicates an origin area which is in Africa only. After finding whether certain cranial or genetic variables are indicative of the expansion from Africa, it was ascertained whether peak points (found in the current analyses and previous studies) were clustered in any region of Africa. A centroid (i.e., as an overall estimate of the origin) was determined from peak points. Some potential caveats were addressed in further analyses centring on i) non-African ancestry, and ii) linearity.

## Method

Analyses of data, and calculations more generally, were through R Versions 4.0.3 (R Core Team, 2020) and 4.0.5 (R Core Team, 2021), and Microsoft Excel. ^12^ BICs were calculated through R or through Microsoft Excel using a formula in Masson (2011) which allows for the BIC to be calculated using *R*^2^ in the formula. When utilising the formula, *R*^2^ was not always employed in the present research. For instance, sometimes squared semi-partial Pearson correlation coefficients were used instead of *R*^2^, as were squared Spearman correlation coefficients and squared semi-partial Spearman coefficients.

### Cranial

The cranial size dimorphism of Holocene modern humans (and dimorphism within cranial dimensions), which had been calculated from measurements in the Howells (1973, 1989, 1995, 1996) dataset (Cenac, 2022), was used in the present analyses. Measurements (56 dimensions) were from 26 populations (*n*_males_ = 1,256; *n*_females_ = 1,156) (Howells, 1989, 1996), and the calculation of size dimorphism (from geometric means) had utilised a natural log ratio in Smith (1999), i.e., ln[(male size)/(female size)] in the prior study (Cenac, 2022). The cranial size dimorphism (calculated using the Howells data) had previously been adjusted for absolute latitude (Cenac, 2022), and that adjusted dimorphism was employed in the present research.

Analyses of cranial diversity also featured the Howells (1973, 1989, 1995, 1996) data (so did Cenac, 2022). Specifically, male crania from 28 populations (*n* = 1,348) and female crania from the 26 populations (Howells, 1989, 1996) were used and the cranial dimensions were the same 56 used in the dimorphism analysis of Cenac (2022). Cranial form diversity can be quantified by i) standardising each dimension by way of a *z*-score transformation, ii) calculating the variance of each of the standardised dimensions for each population, and iii) taking a mean from the variances of all those standardised dimensions (i.e., a mean variance) per population (e.g., Betti et al., 2009). In the present analyses, this method was used on female crania from the Howells dataset. Through that method, the mean form diversity of male crania and the mean shape diversity of female crania had been calculated beforehand from the Howells data (Cenac, 2022) – these diversities were used in the current analyses. In a previous study, cranial shape diversity was determined amongst male crania in the Howells data (28 populations) by calculating the geometric mean of each cranium, dividing the cranial measurements within a cranium by the geometric mean, and then calculating the mean variances after standardising dimensions (von Cramon-Taubadel & Lycett, 2008). For the current analyses, this technique was used to calculate cranial shape diversity with the same data as that previous study (i.e., male crania in the Howells data). From preceding research, the coefficient (*r*) for the correlation between distance from Africa and the cranial shape diversity of females in the Howells data (Cenac, 2022) was used. Cenac (2022) used the Howells dataset obtained through http://web.utk.edu/~auerbach/HOWL.htm, and so did the present research.^13^

### Genetic

The mitochondrial diversity of 51 populations, featured in the supplementary materials of Balloux et al. (2009), was utilised. Mitochondrial diversity was calculated in Balloux et al. (2009) as the averaged between-person dissimilarity (with each dissimilarity found for a pair of people).^14^ Balloux et al. (2009) sourced genetic data from GenBank and (citing Ingman & Gyllensten, 2006) the Human Mitochondrial Genome Database (mtDB); for details on either GenBank or the mtDB, see Benson et al. (2013) and Ingman and Gyllensten (2006) respectively. The mitochondrial diversity was from the complete mitochondrial genome minus hypervariable segments I and II (Balloux et al., 2009). Additionally, Balloux et al. (2009) did present mitochondrial diversity from hypervariable segment I (of 109 populations in HVRBase++). In the present analyses, for representativeness of mitochondrial diversity in general, the diversity from the complete genome (minus the two segments) was used rather than diversity from that hypervariable segment. Diversity (expected heterozygosity) was found in the supplementary material of Balloux et al. (2009) for X- and Y-chromosomal microsatellites, and autosomal SNP haplotypes of 51 populations, and these diversities were used in the present research. Using the HGDP-CEPH Human Genome Diversity Cell Line Panel, on which information can be found, for instance, in Cann et al. (2002) and Rosenberg et al. (2002), Balloux et al. (2009) calculated X- and Y-chromosomal microsatellite heterozygosities. Balloux et al. (2009) found SNP haplotype heterozygosity in Li et al. (2008); Li et al. (2008) calculated mean heterozygosity from HGDP-CEPH data. From summing population sample sizes in Balloux et al. (2009), *N* is 633 for mitochondrial and 963 for autosomal and regarding the allosomes.

### Geographical distances

As for geographical distances, ones calculated beforehand (Cenac, 2022) were used in the present research. Also used were additional distances which were calculated through a means featured by Williams (2011) on http://edwilliams.org/avform.htm. Coordinates for locations in Africa (Betti et al., 2013), coordinates offered for populations regarding the genetic data (Balloux et al., 2009), and coordinates presented elsewhere, including waypoints (Cenac, 2022; von Cramon-Taubadel & Lycett, 2008), were used in the geographical distance calculations. Previous research has used waypoints when calculating geographical distances (e.g., von Cramon-Taubadel & Lycett, 2008) – waypoints were used to calculate distances in the present research. Regarding the waypoints, four of five waypoints listed in a previous study (von Cramon-Taubadel & Lycett, 2008) were featured in calculations like (with the finalised distances) in Cenac (2022); an alternative waypoint, as in Cenac (2022), was made use of rather than the Cairo waypoint which von Cramon-Taubadel and Lycett (2008) used. This alternative waypoint shares the latitude of that Cairo waypoint, but is further east (Cenac, 2022). A distance of 0 km was used for the distance between a population and itself.

Geographical maps (Bartholomew illustrated reference atlas of the world, 1985), including one with representations of locations for populations in the HGDP-CEPH data (López Herráez et al., 2009), were used regarding the calculation of geographical distances. Maps (Bartholomew illustrated reference atlas of the world, 1985) were also utilised for determining which country (or countries) or region peak points, origin areas, and centroids are located in.^15^

### Origins

A number of *origin* coordinates inside/outside of Africa (Betti et al., 2013; von Cramon-Taubadel & Lycett, 2008) were used in the present research. Coordinate pairs were used, in previous research, as *origins* in Africa – 99 in Betti et al. (2013) and three in von Cramon-Taubadel and Lycett (2008) for instance; the present research utilised those 99 coordinate pairs in Betti et al. (2013) and one of the three (Botswana) used by von Cramon-Taubadel and Lycett (2008) as origins too. In von Cramon-Taubadel and Lycett (2008), three origins outside of Africa were used, and these three featured as origins in the present research. Coordinates applied to populations in the Howells data (von Cramon-Taubadel & Lycett, 2008) were also employed in the present research as origins like in Cenac (2022). A number of origin coordinates (Betti et al., 2013) are shown/stated in the present research in figures/text (e.g., the note of Figure 1, Figure S5B).

## Correlation

Correlation tests in the present analyses were two-tailed. When seeing if a variable is associated with geographical distance, previous studies (e.g., Atkinson, 2011; Betti et al., 2013) have employed different methods for choosing which geographical location the population distances are from. For instance, Betti et al. (2013) used the centroid of the origin area. Atkinson (2011) utilised the lowest BIC; in a number of analyses in the present research, the lowest BIC was used. Indeed, if the lowest BIC was found to be at a location outside of Africa for a diversity (and the location gave a negative *r* correlation coefficient), a correlation test (through the cor.test R function) was to be run between diversity and the distance from the origin which gave that lowest BIC. Like previously (Cenac, 2022), Botswana functioned as the origin in Africa for correlation tests regarding the cranium.

### Origin area

As in earlier research (e.g., Betti et al., 2009; Manica et al., 2007), the BIC was the metric for finding which geographical origin gives the peak relationship between variables, and for deciding on the other geographical origins which are just as good as the peak. For each variable, the origin giving the lowest (and therefore strongest) BIC was found, and locations within four BICs of this value were defined as being equally as good a fit (e.g., Manica et al., 2007) for the relationship of geographical distance and the variables (diversities and unadjusted/adjusted dimorphism). However, as alluded to in the *Introduction*, if there were competing geographical areas of best fit for a variable, correlation coefficients were considered when defining the origin area.

Betti et al. (2013) used 99 coordinate pairs alongside others to present two maps of Africa and beyond. Each showed coordinates included and not included in the origin area and the peak point regarding the hip bone shape diversity of males (one graph) and females (the other graph). Following Betti et al. (2013) (who used origins within and outside of Africa), 99 coordinate pairs (found in Betti et al., 2013) were used as origins in Africa, and were employed to display maps of Africa, with it being indicated whether origins were, or were not, in the estimated origin area (figures with these maps indicating inclusion in the origin area, or not being included, are in Table S1). Moreover, in the present research, those 99 coordinate pairs were used to make maps of Africa that had peak points (figures featuring the maps with the peak points are listed in Table S1). Regarding the 99 *origin* coordinate pairs (in Africa) from Betti et al. (2013) which were used in the present research – Betti et al. (2013) utilised coordinates at 5° intervals, e.g., (30°S, 30°E), (25°S, 30°E) etc.^16^

### Graphs

The symbols used for the continents of populations/origins (e.g., Figure 1B) continued, or were adapted, from symbols used in Cenac (2022). Indeed, graphs have extents of similarity to ones in Cenac (2022) in terms of design/axes/labelling, for instance, BIC graphs (e.g., Figure 1B) and correlation coefficient graphs (e.g., Figure S2).^17^ Hunley et al. (2012) presented correlation coefficients for diversity and distance (on the *y*-axis) against geographical regions (*x*-axis).

Geographical regions consisted of locations used as origins (but with one *origin* location being used in the Australia category) (Hunley et al., 2012). Cenac (2022), similarly, displayed correlation coefficients too (*y*-axis), however, these coefficients were plotted against the geographical distance between origins and some location (*x*-axis). For certain graphs (figures stated in Table S1), following Cenac (2022), associations between distance and a variable were plotted on a graph (or, unlike Cenac, 2022, BICs were represented) when each one of 32 geographical coordinate pairs was utilised as an origin. The same *origin* locations (32) used in Cenac (2022) were featured in these graphs. Those 32 coordinate pairs used in Cenac (2022) were from von Cramon-Taubadel and Lycett (2008) for Beijing, Delhi, Tel Aviv, 28 populations in the Howells dataset, and Botswana. The Botswana coordinates are in the north of Botswana (Cenac, 2022). Cenac (2022) constructed graphs using *r*-, *sr*-, and *sr*_s_-values; the present research used *r*-values, but also BICs.

### Positive spatial autocorrelation

Regarding spatial autocorrelation, the three-step method can be used in a Microsoft Excel spreadsheet to find a spatial Durbin-Watson (spatial *DW*) (Chen, 2016) upon which the Durbin-Watson bounds (Savin & White, 1977) can be utilised (Chen, 2016). This method, through the spreadsheet, and bounds were used to check for positive spatial autocorrelation in the correlation analyses regarding the cranium and the Y chromosome. Bounds at the 5% significance level (Savin & White, 1977) were looked to. Within the Chen (2016) spreadsheet is a graph. In any instance of positive spatial autocorrelation, or uncertainty with regard to if such autocorrelation occurred, the graph (i.e., with data in the current analyses put into the spreadsheet) was examined to see which datapoints may be the cause.

### Origin centroid

Locations which lead to the strongest relationships numerically (out of locations in Africa) between geographical distance and diversity/dimorphism (using populations within and outside of Africa), whether calculated through *r*-values (Ramachandran et al., 2005) or BICs, were found from previous research or calculated in the present analyses. Also, a location found through bootstrapping (Tishkoff et al., 2009) was included. Locations were with respect to relationships between distance and i) cranial shape diversity (males, females), ii) cranial size dimorphism, iii) mitochondrial diversity (the present research), iv) autosomal diversity of microsatellites regarding heterozygosity (Ramachandran et al., 2005), repeat lengths (Tishkoff et al., 2009), and SNP haplotype heterozygosity, and v) X-chromosomal diversity (present research). These types of diversity/dimorphism each suggest an estimated origin area in Africa alone (*Results and discussion*; Manica et al., 2007; Tishkoff et al., 2009). The ability of cranial form diversity to indicate the expansion from Africa is suggested to arise more from diversity in cranial shape rather than cranial size diversity (*Results and discussion*). Therefore, to reduce redundancy between origin locations, the locations with respect to cranial form diversity were not included when determining the centroid.

Using the geosphere (Version 1.5-10) package (Hijmans, 2019) in R, a midpoint and centroids were calculated: i) a midpoint was taken between the coordinates from cranial shape diversity (as there was one set of coordinates for males, and one for females), ii) a geographical centroid was calculated from the three locations from the autosomal diversities, iii) the midpoint (cranial shape diversity), the centroid (autosomal diversity), and the three other locations (cranial size dimorphism, mitochondrial diversity, and X-chromosomal diversity) were used when calculating the **expansion origin centroid**.

### Sub-Saharan Africa and admixture

Autosomal microsatellite (mean) expected heterozygosity for each of 106 populations in Tishkoff et al. (2009) (Pemberton et al., 2013) was used. In the present analyses, these 106 populations were included as sub-Saharan African based on categories in Tishkoff et al. (2009). Tishkoff et al. (2009) used categories (for populations in Africa) of central, eastern, southern, western, and Saharan. The central, eastern, southern, and western populations were included as sub-Saharan African populations in the current analyses, unlike Saharan (thereby largely corresponding to Figure 2A of Tishkoff et al., 2009). From population sample sizes in Pemberton et al. (2013), *N* = 2340.

**Figure 2.**
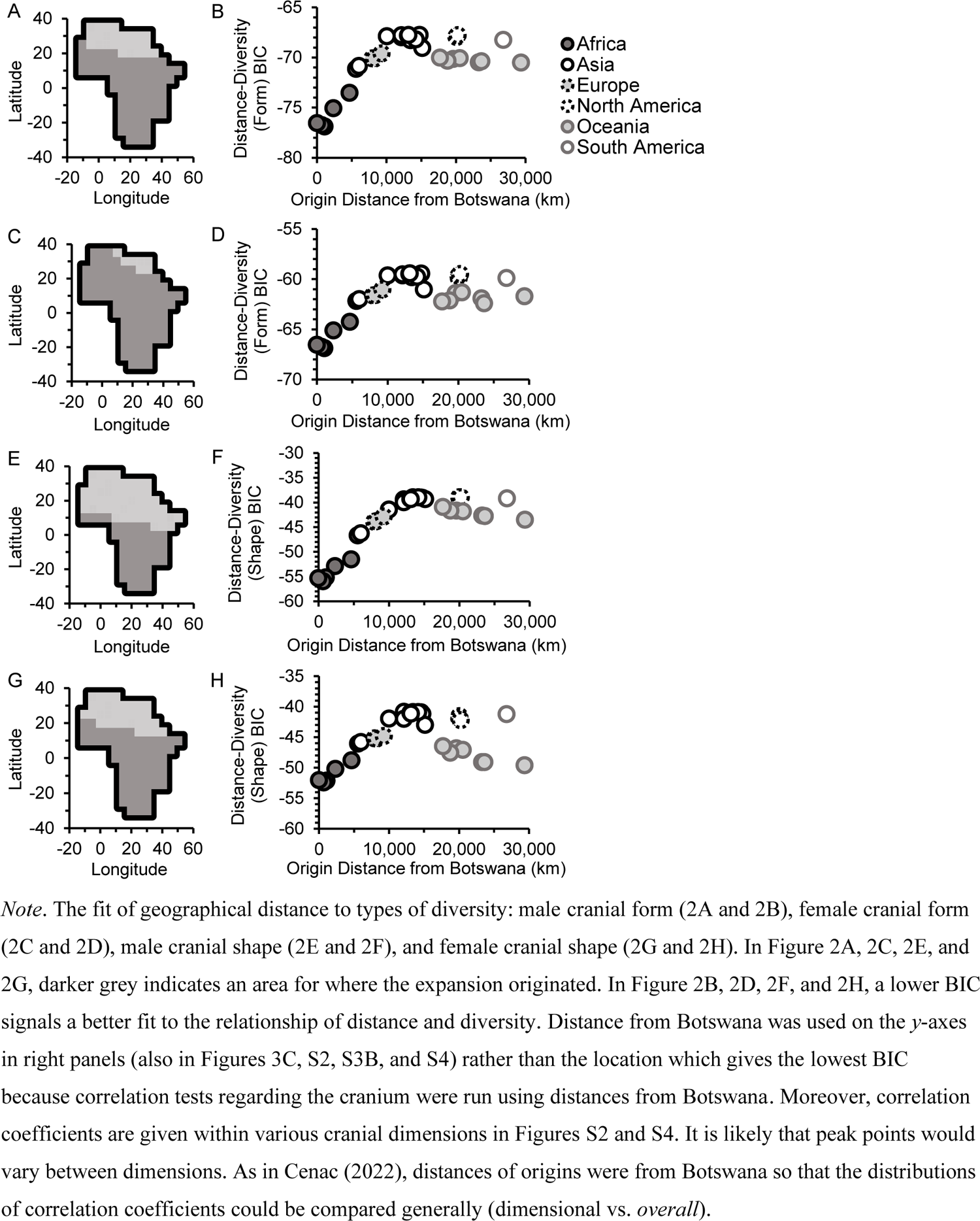
Cranial Diversity Suggests African Origin of Worldwide Expansion

**Figure 3.**
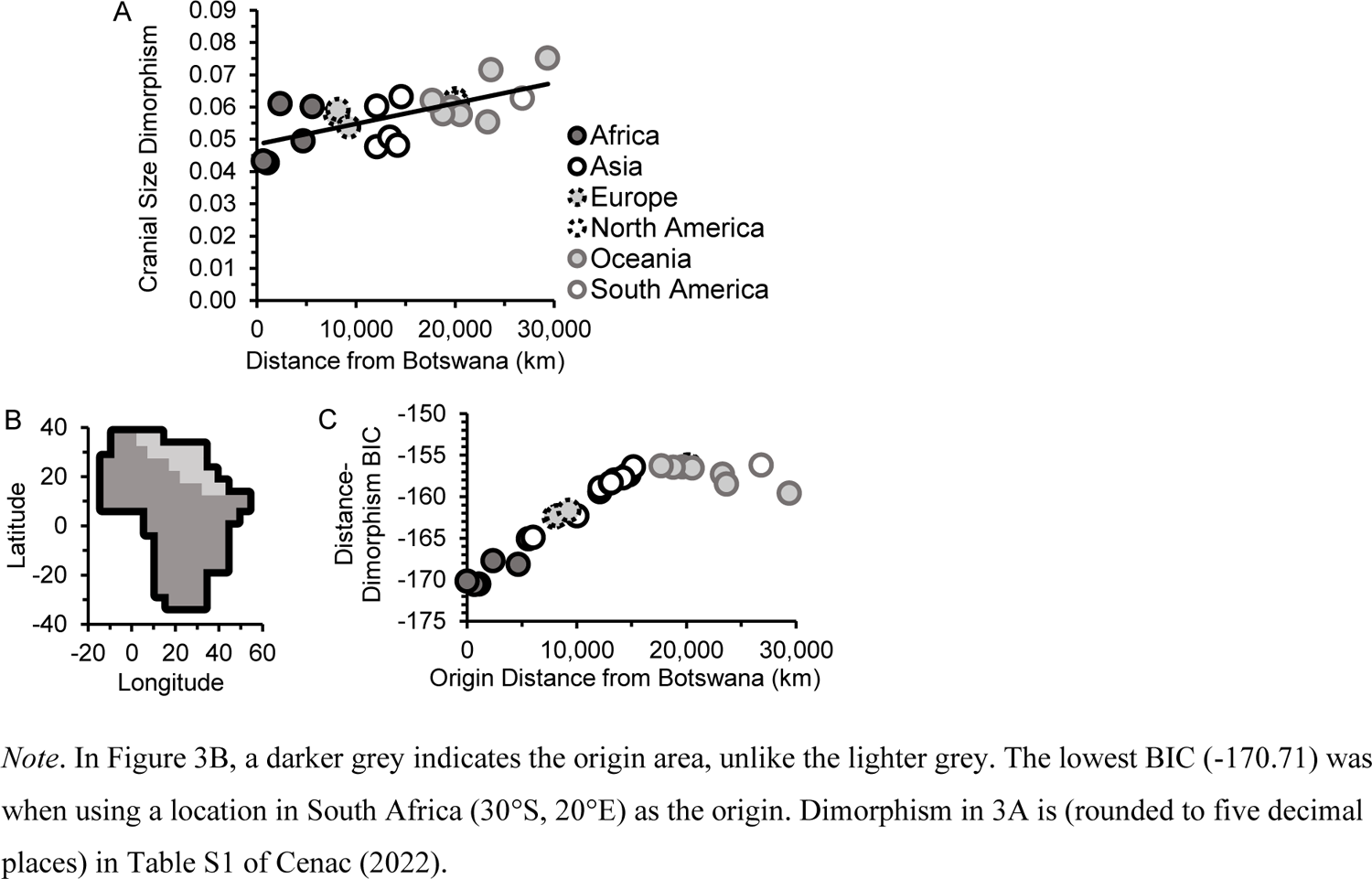
Cranial Size Dimorphism Indicates the Origin of the Expansion

Following some previous research concerning geographical distance and genetic diversity (Pemberton et al., 2013), populations that do not have a population sample size of five or more were not featured in the present analyses. To avoid overlap with Ramachandran et al. (2005), who used autosomal microsatellite heterozygosity of HGDP-CEPH data (the origin centroid in the present research was contributed towards by the peak point in Ramachandran et al., 2005), heterozygosity for the HGDP-CEPH populations was not used despite being available in Pemberton et al. (2013). Estimates of different types of ancestry (East Asian, European / Middle Eastern, Indian, Native American, and Oceanian), as proportions, (Tishkoff et al., 2009) were summed together to represent non-African ancestry.^18^ Parametric and nonparametric analyses were conducted. BICs and correlation coefficients (*r, sr, r*_s_, or *sr*_s_) were found regarding geographical distances (between locations in Africa and populations) and the heterozygosity of the populations which either had or had not been adjusted for non-African ancestry (in nonparametric analysis, a rank-transformation had been applied to heterozygosity, distances, and non-African ancestry). Geographical distances were found as described above, i.e., through Williams (2011). Locations in Africa were from Betti et al. (2013). Coordinates for populations were found in Tishkoff et al. (2009). From the location which had the lowest BIC (alongside a negative correlation coefficient), it was tested whether distance to populations is correlated with the unadjusted/adjusted heterozygosity. For zero-order correlation tests, cor.test in R was used. ppcor (Kim, 2015) was employed in R regarding the tests for semi-partial correlations. It was seen whether autosomal heterozygosity (whether adjusted for non-African ancestry or not) correlates with distance from southern Africa, and if non-African ancestry (adjusted for distance from southern Africa or not) correlates with autosomal heterozygosity. As detailed above, positive spatial autocorrelation was looked for (in parametric analyses) via Chen (2016) and the Savin and White (1977) bounds. To conduct the autocorrelation analyses, interpopulation distances were established through Williams (2011), and 0 km was used i) as the distance when populations had the same coordinates and, ii) for the distance a population was from itself.

### Multiple testing

Statistical significance tests were initially from two sets of analyses which were for two manuscripts; one on the cranium and mitochondrial DNA, and the other on the allosomes and the origin centroid. As a result, the Holm-Bonferroni correction was given to *p*-values (Holm, 1979) for each set separately. Regarding each set, a spreadsheet (Gaetano, 2013) was used to apply the correction. Any *p*-value calculated after the combination of manuscripts (i.e., for the correlation tests involving autosomal microsatellite heterozygosity) was added to the second set. The corrections were not used on the spatial autocorrelation tests.

## Results and discussion

### Origin areas

Using the Bayesian information criterion (BIC) to define the origin area (e.g., Betti et al., 2009; Manica et al., 2007), it was found whether the origin area is exclusively in Africa for various biological measures. A summary of some types of outcomes in the present research and previous studies is presented in Table 2.

**Table 2.**
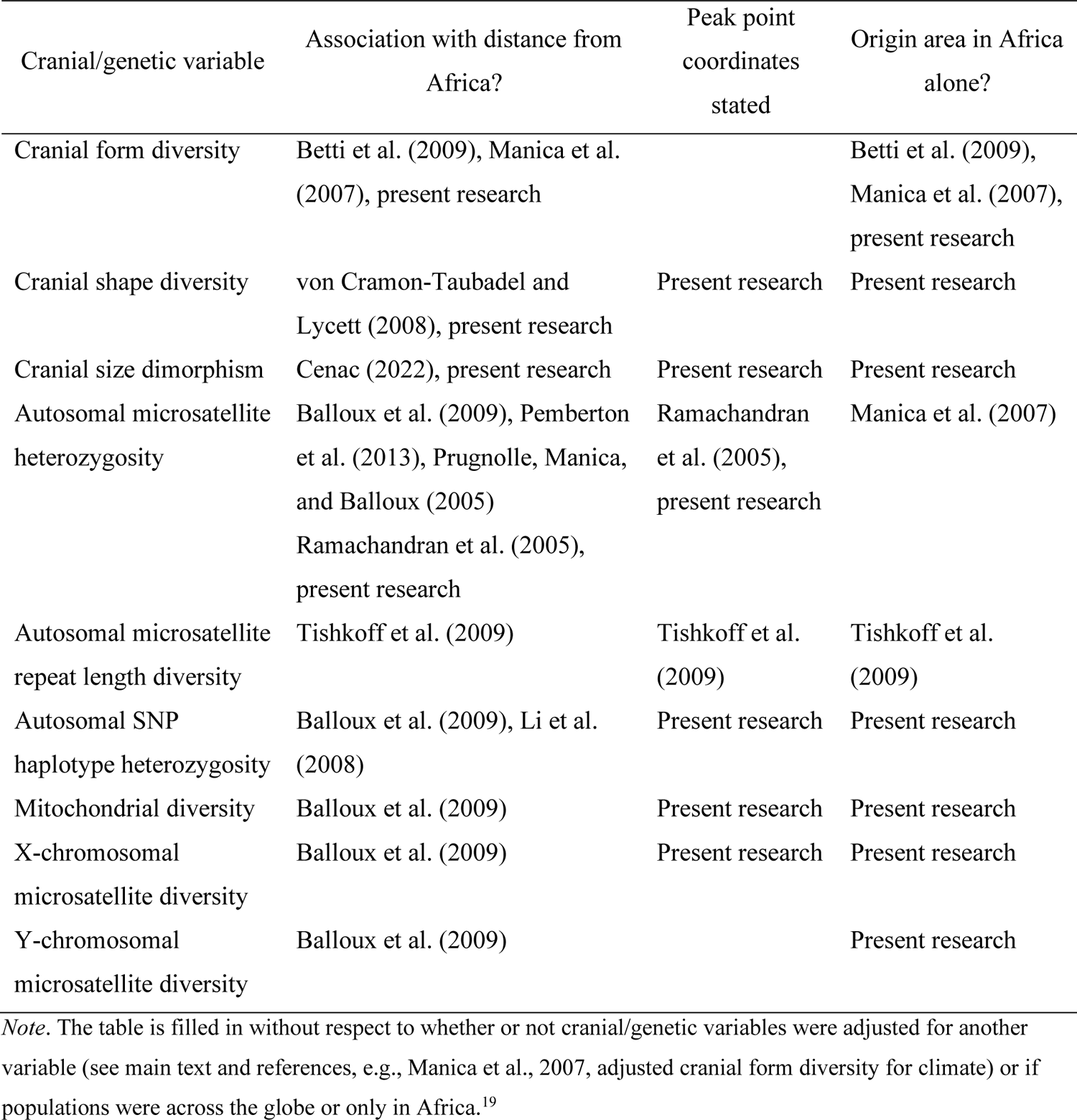
Summary.

| Cranial/genetic variable | Association with distance from Africa? | Peak point coordinates stated | Origin area in Africa alone? |
| --- | --- | --- | --- |
| Cranial form diversity | Betti et al. (2009), Manica et al. (2007), present research |  | Betti et al. (2009), Manica et al. (2007), present research |
| Cranial shape diversity | von Cramon-Taubadel and Lycett (2008), present research | Present research | Present research |
| Cranial size dimorphism | Cenac (2022), present research | Present research | Present research |
| Autosomal microsatellite heterozygosity | Balloux et al. (2009), Pemberton et al. (2013), Prugnolle, Manica, and Balloux (2005) Ramachandran et al. (2005), present research | Ramachandran et al. (2005), present research | Manica et al. (2007) |
| Autosomal microsatellite repeat length diversity | Tishkoff et al. (2009) | Tishkoff et al. (2009) | Tishkoff et al. (2009) |
| Autosomal SNP haplotype heterozygosity | Balloux et al. (2009), Li et al. (2008) | Present research | Present research |
| Mitochondrial diversity | Balloux et al. (2009) | Present research | Present research |
| X-chromosomal microsatellite diversity | Balloux et al. (2009) | Present research | Present research |
| Y-chromosomal microsatellite diversity | Balloux et al. (2009) |  | Present research |

*Note.* The table is filled in without respect to whether or not cranial/genetic variables were adjusted for another variable (see main text and references, e.g., Manica et al., 2007, adjusted cranial form diversity for climate) or if populations were across the globe or only in Africa.<sup>19</sup>

### Cranial

Aligning with previous research (Betti et al., 2009), there was a negative correlation between distance from Africa and cranial form diversity for males, *r*(26) = −.52, *p* = .023, spatial *DW* = 2.50 (Figure S1A), and females *r*(24) = −.49, *p* = .044, spatial *DW* = 2.10 (Figure S1B). Using the Howells data, prior research had already found that the shape diversity of male crania negatively correlates with distance from Africa (von Cramon-Taubadel & Lycett, 2008).^20^ Amongst female crania, distance from Africa correlated negatively with cranial shape diversity, *r*(24) = −.59, *p* = .009, spatial *DW* = 2.26 (Figure S1C). For cranial form diversity, the geographical area of best fit was only in Africa (Figure 2A-2D). Regarding cranial form diversity, in preceding research the area had been entirely in Africa for males (Betti et al., 2009; Manica et al., 2007), but not for females (Betti et al., 2009).

As for cranial shape diversity, the area was just in Africa for male crania (Figure 2E and 2F). In prior research with male crania, *R*^2^s representing the extent of association between shape diversity and geographical distance were numerically greater when distance was from Africa (Botswana, Democratic Republic of the Congo, or Ethiopia) rather than from Asia (China, India, or Israel) (von Cramon-Taubadel & Lycett, 2008) which is congruent with the present research which found that the area of best fit is in Africa. However, for the cranial shape diversity of females, there was more than one geographical area of best fit, with one being in Africa and at least one other being outside of Africa (Figure 2G and 2H). This finding arose from some locations in Oceania having low BICs. For female crania, correlation coefficients were negative for locations in Africa but positive for locations in Oceania (Figure S2C). Because research agrees with a decline in types of diversity with distance from the expansion origin (Ramachandran et al., 2005; von Cramon-Taubadel & Lycett, 2008), the area with negative coefficients is the estimated area of origin (as discussed in the *Introduction*). Therefore, results with female cranial shape diversity are in step with the expansion being from Africa and not elsewhere. As a result, in the present research, cranial diversity in form and shape lead to the estimations of origin areas which are only in Africa.

Cranial shape diversity appears to indicate a smaller origin area than cranial form diversity does (Figure 2). Hence, cranial shape diversity may be better at indicating the origin compared to cranial form diversity. Such an outcome should not be unexpected. This is because shape, along with size, is part of form (Betti et al., 2010) and cranial size diversity (controlled for absolute latitude) has no correlation with distance from Africa (Cenac, 2022).

Regarding a test of whether cranial size dimorphism correlates with distance from Africa (Botswana), positive spatial autocorrelation was at hand when the analysis used 26 populations, spatial *DW* = 1.17. A graph (produced through the Chen, 2016, spreadsheet) indicated that this was caused by two populations: Ainu and North Japan. The positive spatial autocorrelation was resolved by the absence of those populations, spatial *DW* = 1.87. With the remaining populations, it was found that cranial size dimorphism and distance from Africa have a positive correlation, *r*(22) = .66, *p* = .003 (Figure 3A).

Using cranial size dimorphism, an origin area was suggested within Africa alone (Figure 3B and 3C). Therefore, results seem supportive of cranial size dimorphism having a link to the expansion from Africa, and cranial size dimorphism being a useful indicator of this expansion.^21^ Whilst prior research had explored whether cranial size dimorphism, when adjusted for absolute latitude, does have a correlation with distance from Africa (Cenac, 2022), such an adjustment is likely to be without value (Text S1). Nevertheless, with adjusted cranial size dimorphism, an origin area was also found to fully be in Africa (Figure S3). And so, given that adjusted cranial size dimorphism does correlate with distance from Africa (Cenac, 2022; Text S1), cranial size dimorphism appears to remain as an indicator of the expansion after adjusting it for absolute latitude.

The level of correlation between adjusted cranial size dimorphism and geographical distance does not appear to be a good yardstick of coefficients for correlations between the adjusted dimorphism of individual dimensions and geographical distance (Cenac, 2022). This is apparent with unadjusted dimorphism (i.e., zero-order correlation coefficients) (Figure S4A and S4C). Such an occurrence is present regarding relationships between geographical distance and both cranial form diversity (Cenac, 2022; Figure S2A and S2B) and cranial shape diversity (Figure S2C-S2F).

### Genetic

Figure 1 suggests that the origin area regarding autosomal SNP haplotype heterozygosity is totally in Africa. As for mitochondrial diversity, when using 32 worldwide origins, the lowest BIC was in North America, specifically at the Arikara location which had a BIC of 382.75 (Figure 4A). Origins with BICs within four of Arikara were in North America and Africa (Figure 4A). As with cranial shape diversity, one can note the direction of *r*-values. Such values were negative for Africa and positive for North America (Figure 4B). Hence, the geographical area of best fit in Africa can be considered as the estimated origin area for the same reason as with cranial shape diversity amongst females. Using the four-BIC route (Manica et al., 2007), when therefore not considering origin locations in North America, the geographical area of best fit was only within Africa (Figure 4C).

**Figure 4.**
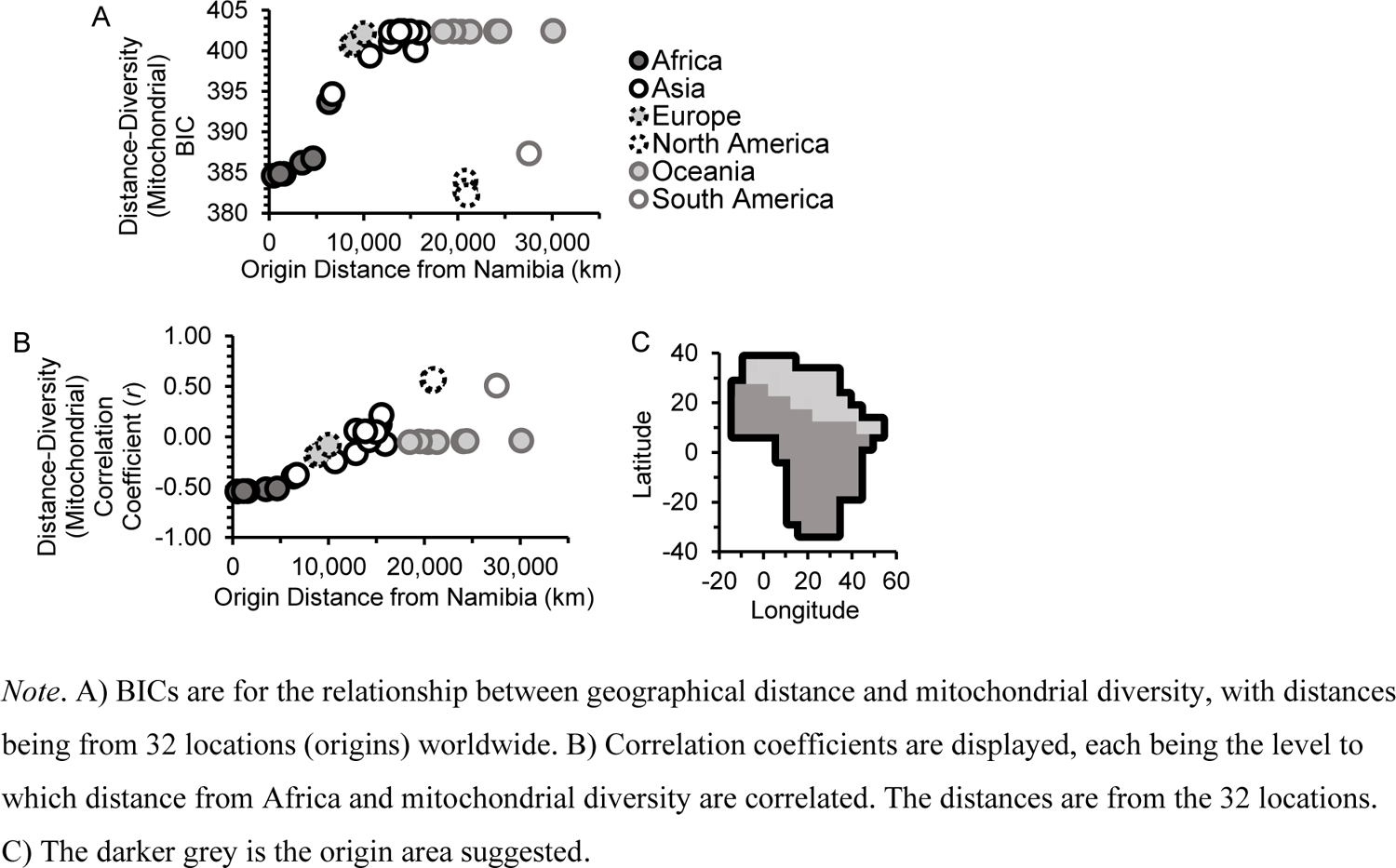
Mitochondrial Diversity Suggests that Africa has the Expansion Origin

Results indicated that the geographical area of best fit for the relationship of X-chromosomal microsatellite heterozygosity and geographical distance was exclusively in Africa (Figure 5A and 5B). Therefore, given that these diversities (autosomal SNP haplotype, mitochondrial, and X-chromosomal) are each associated (negatively) with geographical distance from Africa (Balloux et al., 2009), origin areas support these diversities indeed having an indication of the expansion from Africa.

**Figure 5.**
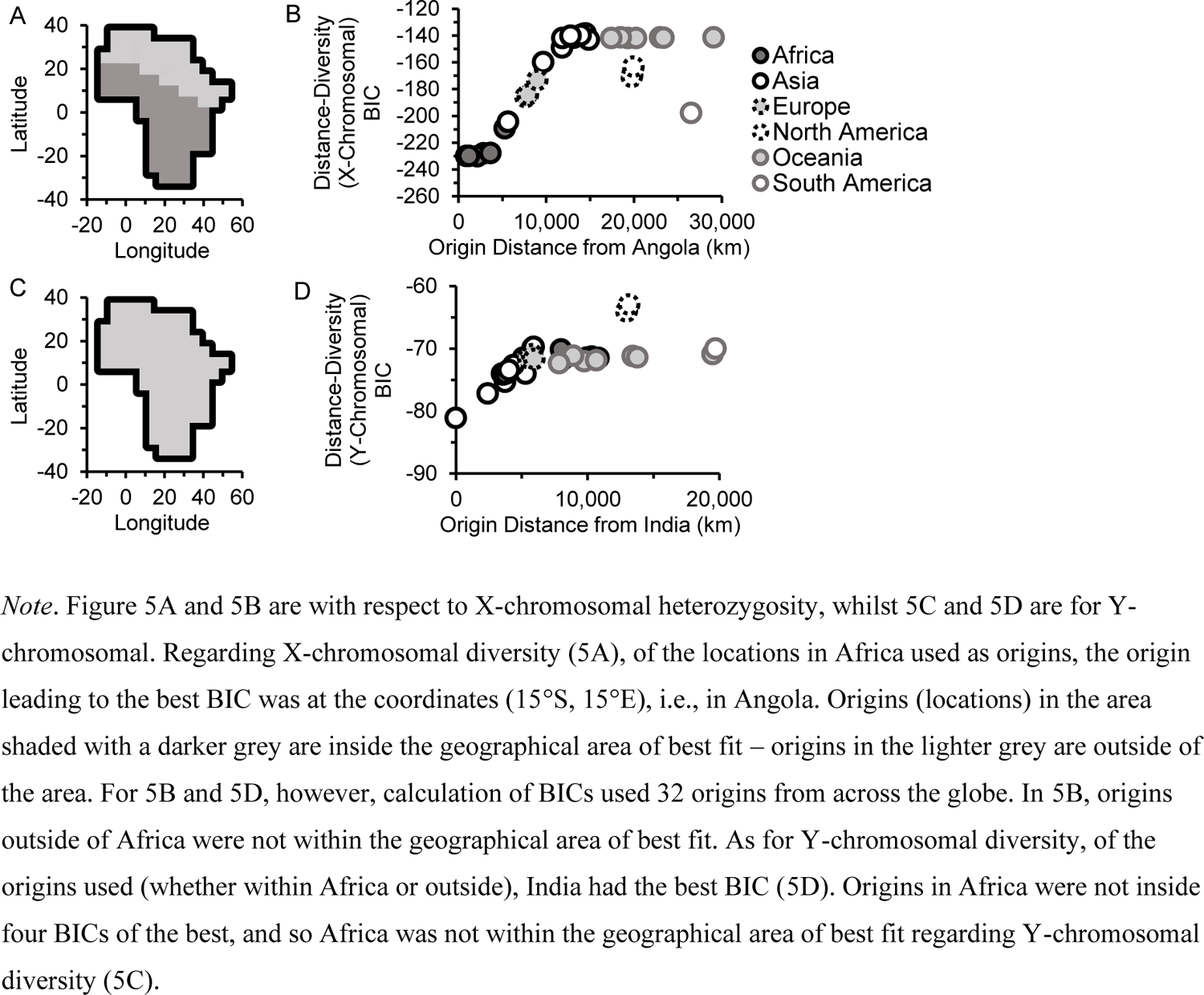
X-Chromosomal, but not Y-Chromosomal, Microsatellite Diversity Signals African Origin for Expansion

With Y-chromosomal microsatellite heterozygosity, the best BIC was found to be within India (Figure 5D). Distance from India did correlate negatively with Y-chromosomal diversity, *r*(49) = −.55, *p* < .001, spatial *DW* = 1.83. All origins in Africa were more than four BICs away from the lowest BIC. Therefore, Africa did not have any of the geographical area of best fit (Figure 5C). No origin (of those used) outside of Asia was within four BICs of the lowest, and so the geographical area of best fit for Y-chromosomal microsatellite heterozygosity was possibly within Asia fully.

However, what if more than one origin was indicated? Previous research had considered this with respect to autosomal heterozygosity in expansion (Manica et al., 2007) and the origin of language (Atkinson, 2011). For instance, Atkinson (2011) constructed models for predicting phonemic diversity from types of geographical distance (distance from the peak point in Africa, distance from elsewhere) and population size. Atkinson (2011) did not find that phonemic diversity (in the model) had a negative correlation with distance from elsewhere. A similar sort of analysis was used in the present research; when adjusting Y-chromosomal diversity for distance from India, the numerically lowest correlation coefficient (*sr*) for the relationship between distance and adjusted diversity when using an African origin was found – distance from this origin was not correlated with the adjusted diversity, *sr*(48) = −.10, *p* = .97, spatial *DW* = 1.83 (the correlation test was run using the ppcor package of Kim, 2015). As a result, although there is a negative relationship between Y-chromosomal microsatellite heterozygosity and distance from Africa (Balloux et al., 2009), this relationship would not appear to actually be reflective of the expansion from Africa.

When using a combined African population and a number of non-African populations, a phylogenetic tree (arising from Y-chromosomal microsatellites) has its root located between Africans and non-Africans (Seielstad et al., 1999), hence Table 1 and Figure 5C and 5D may be surprising.

Nevertheless, studies are congruent with there being migration back into Africa (e.g., Henn, Botigué et al., 2012; Hodgson et al., 2014). Indeed, whilst research with the Y chromosome does suggest expansion from Africa into Asia, migration from Asia to Africa is also indicated (e.g., Hammer et al., 2001). Furthermore, if Y-chromosomal haplogroup E (which has been described as being the most common of Y haplogroups in Africa) was from an expansion from Asia (Wang & Li, 2013), maybe it should not be that surprising if extents of Y-chromosomal diversity do not reflect the worldwide expansion from Africa. Hence, the signal of expansion from Asia in Y-chromosomal microsatellite diversity (Figure 5D) could very well represent migration from Asia to other continents, including back into Africa.

The favouring of expansion from Asia by Y-chromosomal diversity (e.g., Figure 5C and 5D) may, to some degree, align with Hallast et al. (2021). Considering expansion from Africa, one would predict that, outside of Africa, the Y chromosomes of non-Africans would seem to have spread from western Eurasia (Hallast et al., 2021). Nevertheless, in Hallast et al. (2021), outside of Africa, it was indicated that they spread from eastern or southeastern Asia (Hallast et al., 2021). This could be explained by more easterly Y chromosomes having replaced westerly ones (Hallast et al., 2021); replacement could, therefore, have been a factor regarding findings with Y-chromosomal microsatellite diversity in the present research.

Nevertheless, in Balloux et al. (2009), Y-chromosomal diversity was calculated from six microsatellites, whilst X-chromosomal and autosomal microsatellite diversities were determined from 36 and 783 microsatellites respectively. Ten Y-chromosomal microsatellites is few (Seielstad et al., 1999). A tally of six microsatellites could be described similarly. Therefore, the apparent signalling of expansion from Asia (rather than from Africa) in Y-chromosomal diversity (Figure 5C and 5D) was found through few microsatellites. It would be unwise to discount Y-chromosomal microsatellite diversity from reflecting the expansion from Africa until far more microsatellites are used.

Finding genetic diversity to not be greatest in Africa can be indicative of ascertainment bias (e.g., Conrad et al., 2006, with SNP heterozygosity). Previous research has considered if there are indications of ascertainment bias regarding microsatellites in the HGDP-CEPH data (e.g., Ray et al., 2005, who utilised microsatellite data used in Rosenberg et al., 2002, and Rosenberg et al., 2002, employed autosomal microsatellite data) – perhaps future research could address whether ascertainment bias is present in those six Y-chromosomal microsatellites.

### Expansion origin centroid

Because Y-chromosomal diversity was not found to signal the expansion from Africa, coordinates regarding Y-chromosomal diversity were not included when calculating the origin centroid – an overall indication of the launching point of the expansion (Figure 6).

**Figure 6.**
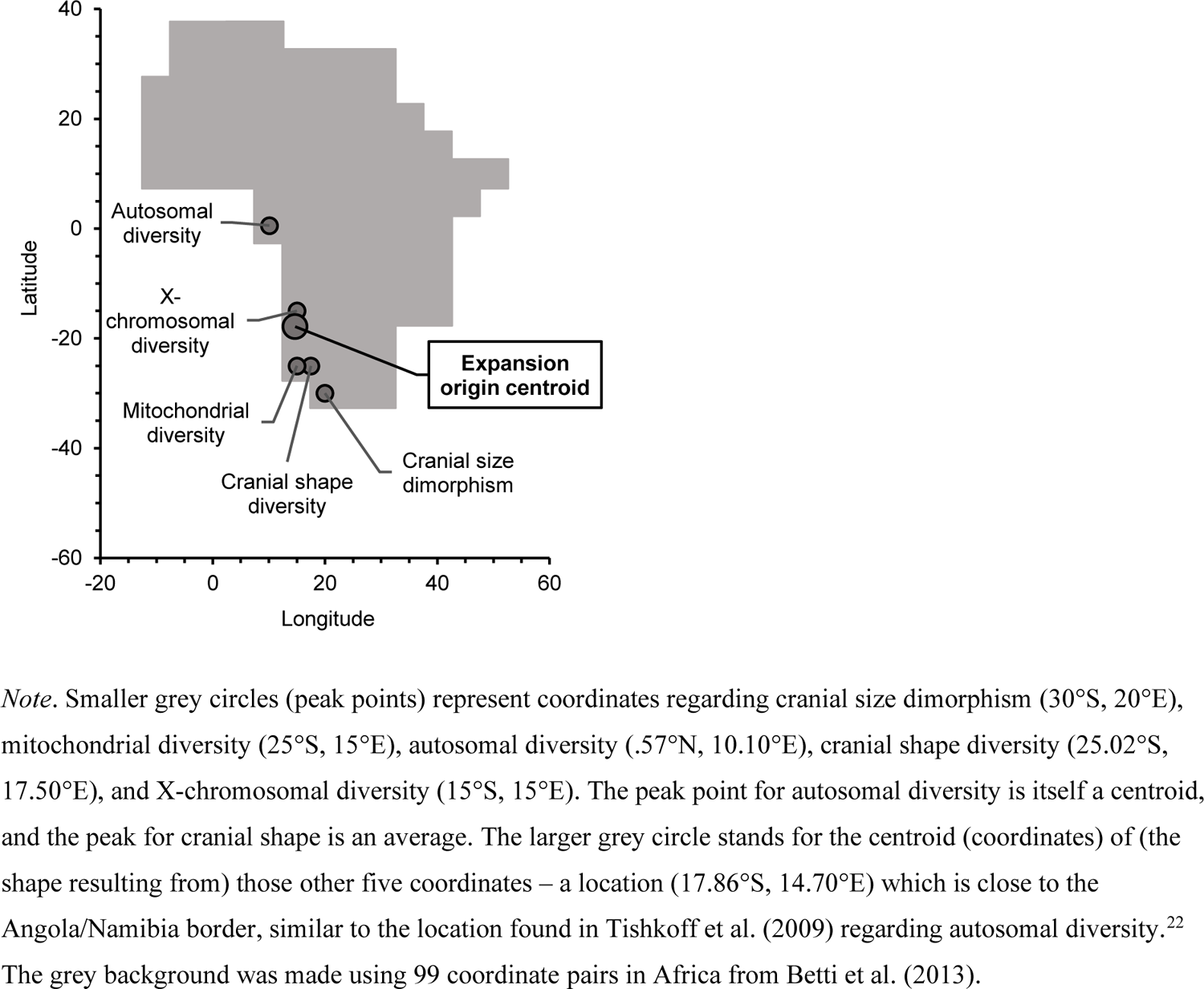
The Expansion Originated in Southern Africa?

Given the lack of support for using an adjustment for absolute latitude on cranial size dimorphism (Text S1), the expansion origin centroid (Figure 6) was not calculated using the adjusted dimorphism. And so, when calculating the centroid, the cranial size dimorphism peak point was used alongside a number of peak points for diversities which were either from the present research or previous research. These diversity peak points were mitochondrial, X-chromosomal, cranial shape diversity (an average of peak points for males and females), and autosomal. The autosomal diversity *peak point* was really a centroid of three peak points. These three peak points were from autosomal microsatellite repeat length diversity (Tishkoff et al., 2009), autosomal microsatellite heterozygosity (Ramachandran et al., 2005), and autosomal SNP haplotype heterozygosity.

The shape used when estimating the expansion origin centroid was formed of lines connecting coordinates (with coordinates being vertices) – from the autosomal diversity coordinate to mitochondrial diversity to cranial size dimorphism to cranial shape diversity to X-chromosomal diversity to autosomal diversity. Regarding indicators of the expansion from Africa, most of the types of peak points are in southern Africa (four out of five), which generally supports southern Africa being the origin (Figure 6). Indeed, a centroid of the peak points suggests that southern Africa is the most probable origin of the expansion (Figure 6), in the context of the peak points generally clustering in the southern African region (like in Choudhury et al., 2021, Angola, Malawi, and Zambia were regarded as part of southern Africa in the present research).

### Number of populations

Going by preceding research/publications (Balloux et al., 2009; Betti et al., 2009; Bartholomew illustrated reference atlas of the world, 1985; Howells, 1989, 1996; López Herráez et al., 2009; Ramachandran et al., 2005; Tishkoff et al., 2009), few populations in Africa were involved when finding peak points except for one of the three types of autosomal diversity – autosomal repeat length diversity.^23^ Given previous research on phonemic diversity (Van Tuyl & Pereltsvaig, 2012), a concern may be the instances in which peak points were calculated from populations globally but only including a small number of populations in Africa. When considering the relationships between geographical distance and a variable more generally than Van Tuyl and Pereltsvaig (2012), it seems that there is variety in how close peak points in Africa are when the peaks are found with populations worldwide compared to populations in Africa (Figure 7).

**Figure 7.**
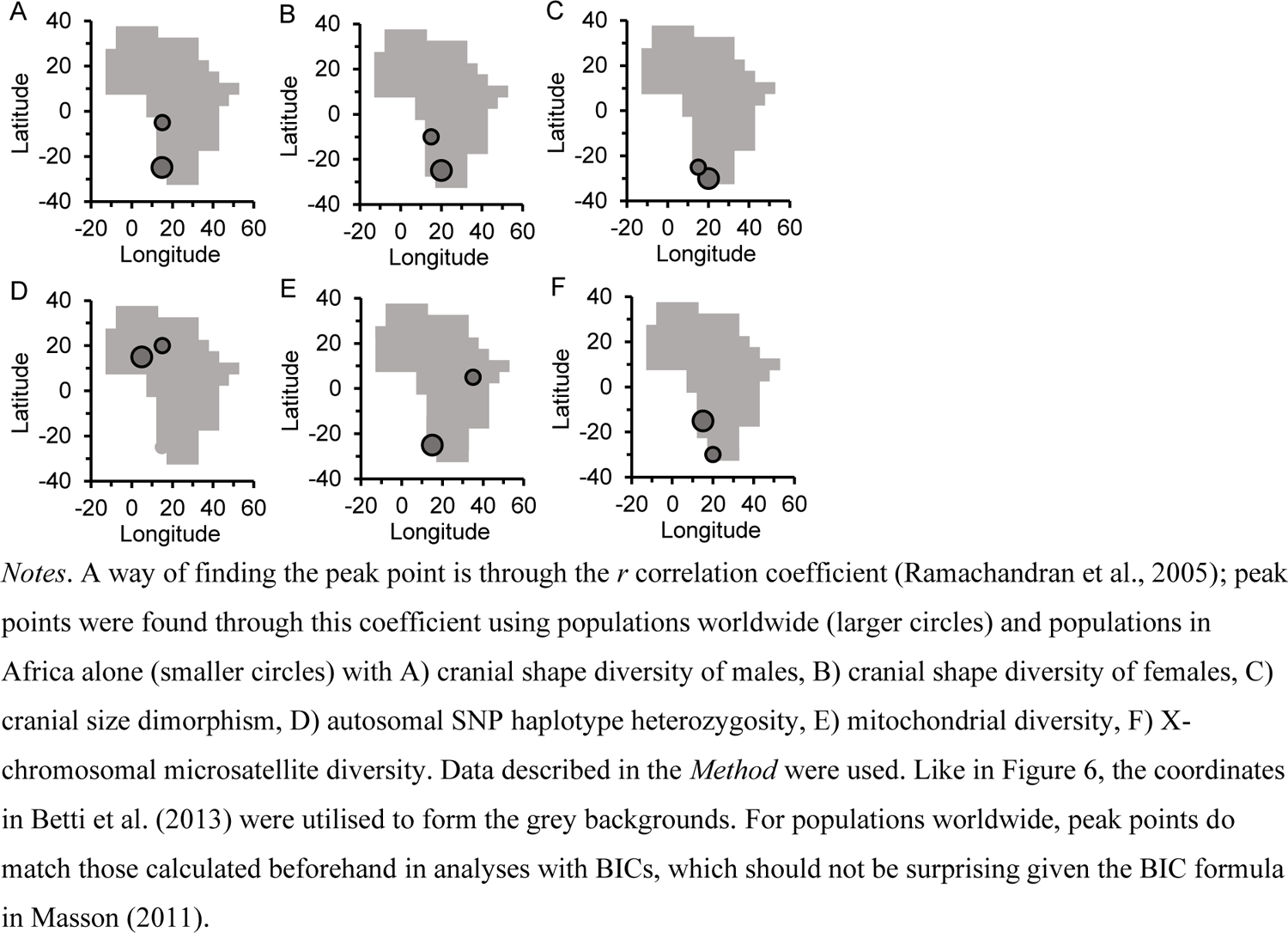
Peak Points with Populations Globally, and in Africa

Yet, Figure 7 suggests there indeed appears to generally be agreement between peak points regardless of whether they are calculated from populations worldwide or in Africa only. And so, peak points in Africa (when using populations across the globe) are not defined exclusively by African populations, but may only be particularly *driven* by such populations (cf. Van Tuyl & Pereltsvaig, 2012). Therefore, if the distribution of biological diversity/dimorphism in Africa drives where the peak point in Africa is found to be, there could indeed be a concern over the number of populations in Africa that were featured when calculating the origin centroid in Figure 6. However, the origin centroid did arise from different types of diversity/dimorphism variables (and some different samples – see *Method*), which should counter the issue of small numbers of populations to an extent.

### Diversity within Africa and non-African ancestry

Using populations across the world, autosomal microsatellite heterozygosity i) is found to negatively correlate with distance from Africa (Prugnolle, Manica, & Balloux, 2005), and ii) results in an origin area that is fully in Africa (Manica et al., 2007). If the location of the origin centroid *within* Africa is greatly due to populations in Africa, one would assume that the signal of the expansion is clear amongst populations in Africa (although correlations between distance and autosomal diversity inside Africa were not present in Hunley & Cabana, 2016, but see proceeding paragraphs).

In the present research, the autosomal microsatellite heterozygosity in 106 sub-Saharan African populations (calculated from Tishkoff et al., 2009, data by Pemberton et al., 2013) was used to indicate if there is a signal of the expansion within Africa. The magnitude of relationships between autosomal microsatellite heterozygosity in sub-Saharan Africa and geographical distance (within Africa) was strongest when distance was from Angola (15°S, 20°E), with there being a negative correlation between heterozygosity and distance from that location, *r*(104) = −.31, *p* = .006, spatial *DW* = 2.01, and an origin area in southern Africa was indicated (Figure 8A).^24^ (This is by no means the first analysis to explore whether distance and a variable are associated within Africa, e.g., Henn, Gignoux et al., 2011, did explore this.) Therefore, autosomal microsatellite heterozygosity within sub-Saharan Africa seems to indicate a southern African origin of an expansion within Africa. Assuming that this expansion is part of the worldwide expansion, then the origin of the worldwide expansion appears to be indicated. Hence, this origin indicated with populations in Africa (Figure 8A) broadly agrees with the origin centroid.

**Figure 8.**
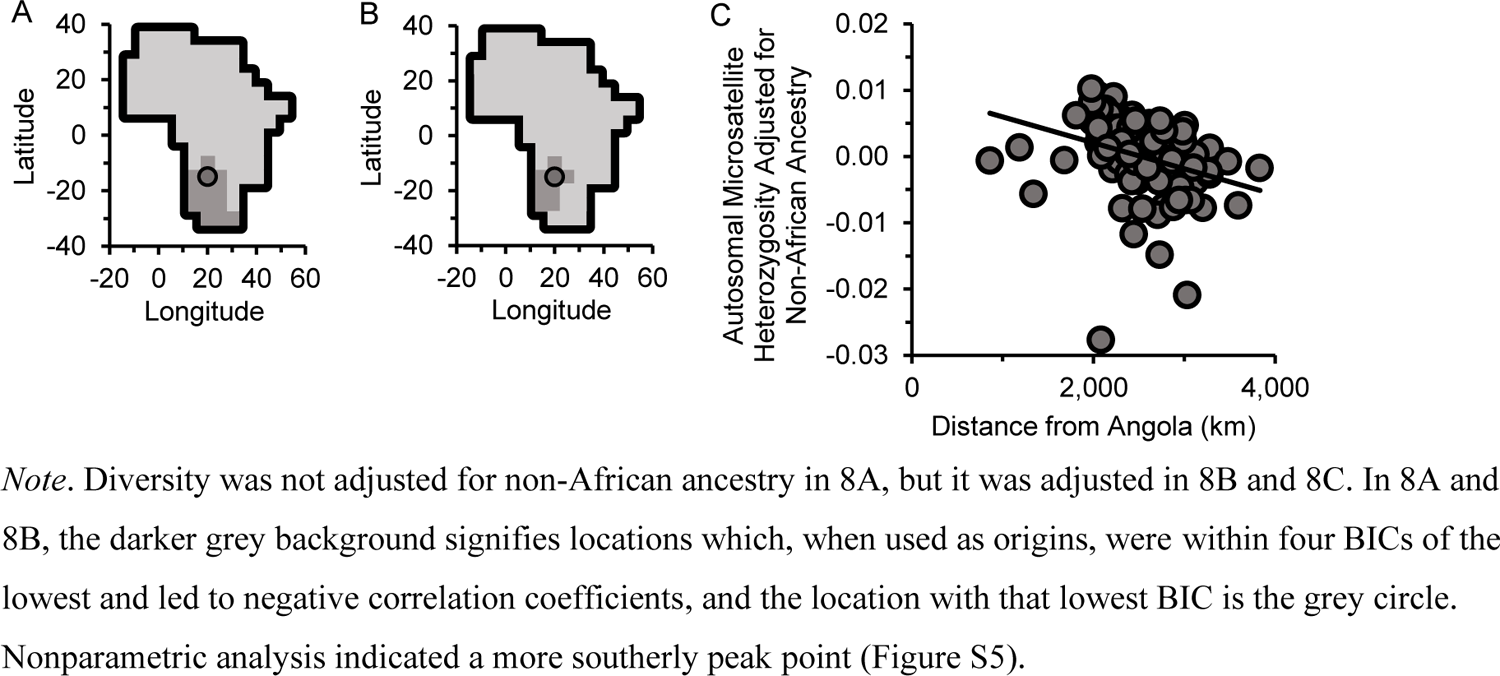
Origin of the Expansion from the Autosomal Microsatellite Diversity of Populations in Sub-Saharan Africa

However, a graph placing distance (from the peak) against microsatellite heterozygosity suggested that heteroscedasticity (i.e., non-constant variability) was present. Therefore, the confintr package (Mayer, 2022) was used in R to bootstrap a 99% confidence interval regarding the correlation coefficient (*r*) for the relationship between distance from the peak and heterozygosity. The default method of bias-corrected accelerated (Mayer, 2022) was chosen. In bootstrapping, a total of 1,000 samples was used – to match the number of bootstrap samples employed when comparing correlation coefficients to each other in the analysis presented in Text S1. The confidence interval [-.51, −.01] did not include 0 and it consisted entirely of negative values, thereby favouring there being a negative correlation. Furthermore, nonparametric analysis supported the presence of a negative correlation, albeit with a different peak point which was (25°S, 15°E), *r*_s_(104) = −.51, *p* < .001 (Figure S5A), with the origin area still being indicated to fully be in southern Africa (Figure S5B).

Nevertheless, Hunley and Cabana (2016) did not find a correlation between distance from eastern Africa and autosomal microsatellite diversity within any of three categories of populations in Africa. Yet an absence of correlations should not be surprising because, in the present research, eastern Africa was not in the origin area for the relationship between distance and heterozygosity in sub-Saharan Africa (Figure 8A), with southern Africa looking to be a more plausible origin for the expansion than eastern Africa (Figures 6 and 8A).

### Non-African ancestry

Keeping in mind previous research with respect to admixture (Gallego Llorente et al., 2015; Haber et al., 2016; Pickrell et al., 2014; Pickrell & Reich, 2014), non-African ancestry may underlie support for a southern African origin. For instance, Pickrell et al. (2014) presented extents of West Eurasian ancestry for a number of populations. An observation of theirs (Pickrell et al., 2014, p. 2636) certainly can be interpreted as referring to West Eurasian ancestry in populations largely being lower in southern Africa than in eastern Africa. Moreover, other research is supportive of West Eurasian ancestry being greater in eastern Africa than in southern (and western) Africa (Gallego Llorente et al., 2015). Therefore, perhaps West Eurasian (and more generally non-African) ancestry contributed towards the apparent support in the present research for the origin of a worldwide expansion being in southern Africa.

The heterozygosities found from the Tishkoff et al. (2009) data (Pemberton et al., 2013) were used to examine whether the expansion origin is indicated to be in southern Africa when autosomal heterozygosity in sub-Saharan Africa is adjusted for non-African ancestry (distance was not adjusted).^25^ With the adjusted heterozygosity, an origin area in southern Africa was found (Figure 8B). The location at which the BIC was lowest was the same as with unadjusted heterozygosity (parametric analysis). Distance from (15°S, 20°E) was negatively correlated with adjusted heterozygosity, *sr*(103) = −.32, *p* = .008, spatial *DW* = 2.00 (Figure 8C). However, heteroscedasticity was indicated to occur (Figure 8C). Therefore, ppcor (Kim, 2015) was used to run a semi-partial Spearman correlation between distance from the peak of (25°S, 15°E) and autosomal diversity when the diversity was adjusted for non-African ancestry; there was a correlation, *sr*_s_(103) = −.54, *p* < .001 (Figure S5C). Moreover, southern Africa was suggested to have the expansion origin area (Figure S5D). Therefore, the signalling of a southern African origin through autosomal heterozygosity would not appear to be attributable to non-African ancestry.^26^

Results suggest that autosomal microsatellite heterozygosity within sub-Saharan Africa reflects part of the worldwide expansion. This finding is favourable towards the origin centroid (Figure 6) actually approximating where the worldwide expansion started. Nonetheless, it is not clear if an expansion signal in autosomal microsatellite heterozygosity is present in regions outside of Africa – if the signal is not in other regions (Text S2), it seems questionable whether the signal in Africa truly reflects the expansion. Given populations and their locations in Tishkoff et al. (2009), the present research used relatively few populations inside southern Africa (four populations) in the analyses with sub-Saharan African heterozygosity. Indeed, not many populations were close to the location of the lowest BIC (e.g., Figure 8C). This does undermine some confidence in whether adjusted/unadjusted heterozygosity actually declines linearly in sub-Saharan Africa as distance from southern Africa increases (see the literature regarding expansion and linearity, e.g., Deshpande et al., 2009, Hunley & Cabana, 2016, Liu et al., 2006, Prugnolle, Manica, & Balloux, 2005).

### Linearity

Indeed, Figure 8C may hint at there being a non-linear (and non-monotonic) trend. Monotonic associations had been measured in the present research – building on reasoning in Liu et al. (2006), a lack of consideration for non-monotonic trends may present an issue for estimating where the expansion originated (Text S3). Further analysis, upon the removal of two atypical datapoints, supports there being a non-monotonic trend (a quadratic trend) whether or not heterozygosity is adjusted for non-African ancestry (Text S3). Moreover, an origin area only in southern Africa is indicated amongst quadratic models (Figures 9 and 10).

**Figure 9.**
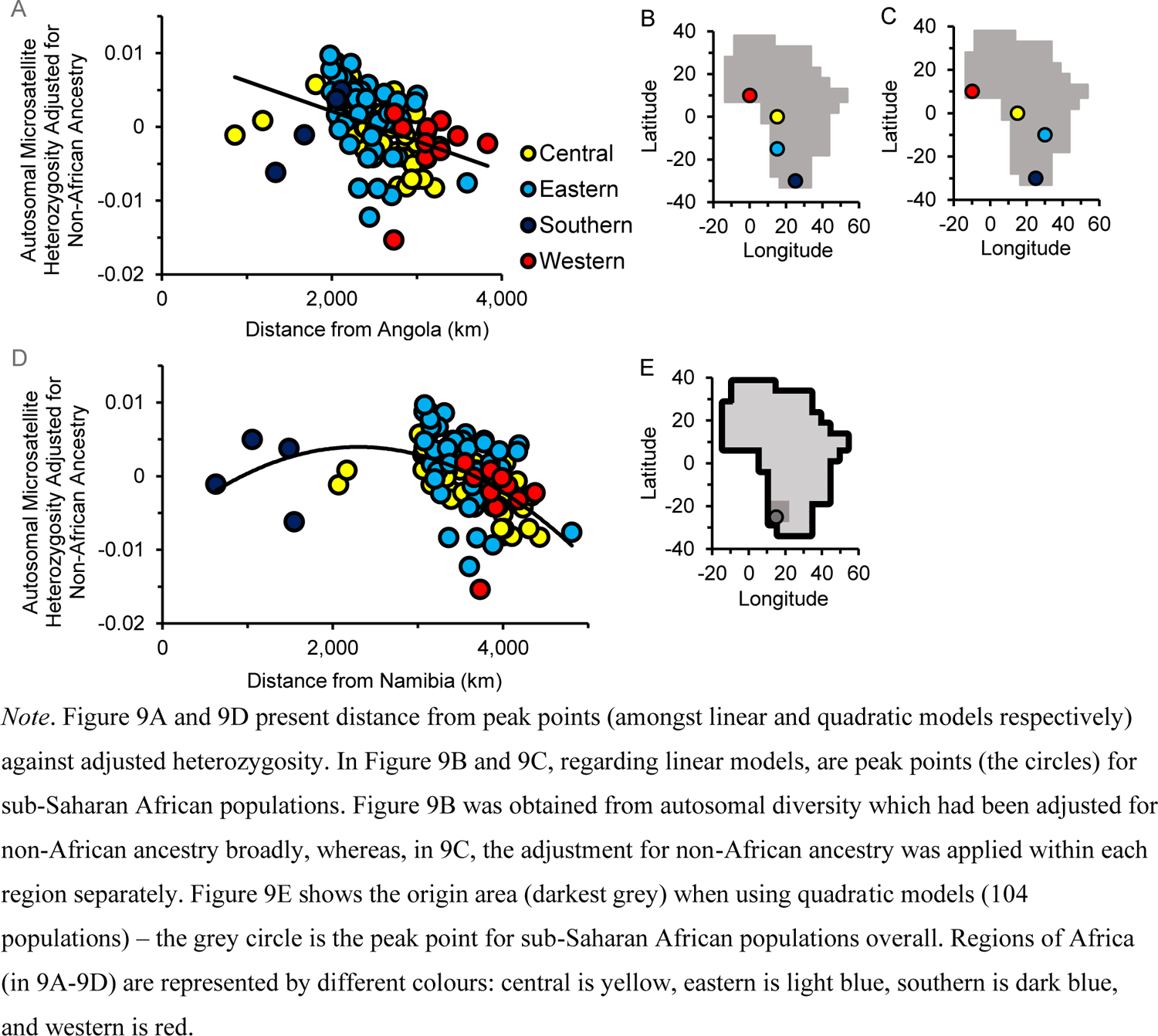
Autosomal Microsatellite Diversity: Adjusting for Non-African Ancestry

**Figure 10.**
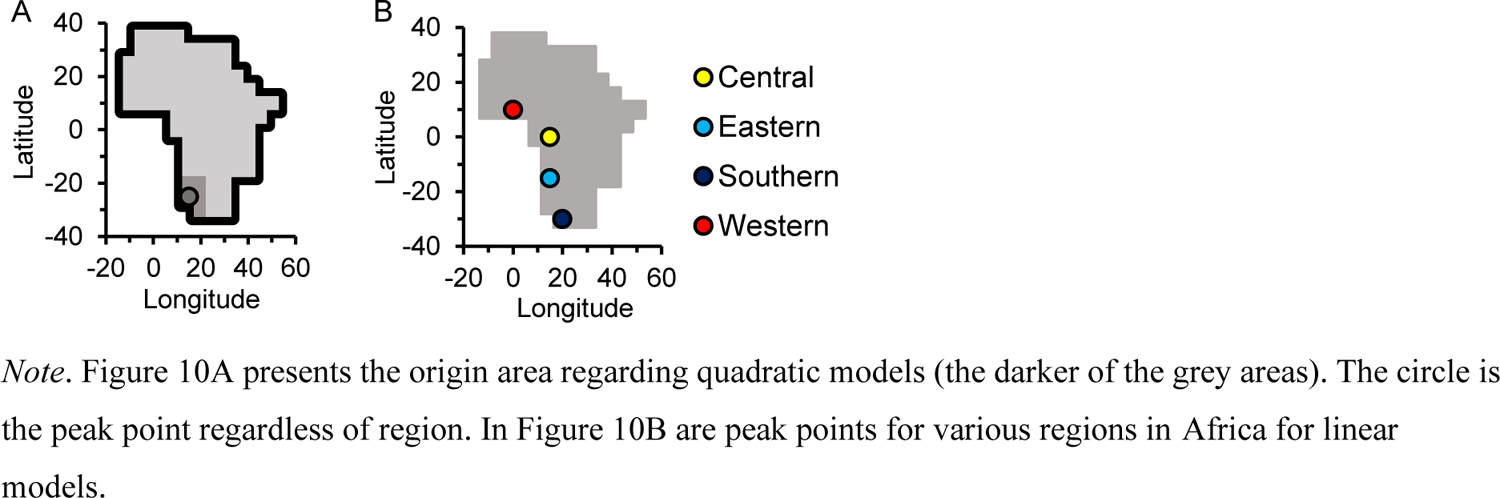
Autosomal Microsatellite Diversity: Not Adjusted for Non-African Ancestry

With adjusted (Figure 9D) and unadjusted heterozygosity, there was possibly an initial increase of diversity (to circa. 2,000-3,000 km) and then a decline.^27^ This possible increase then decline is reminiscent of Figure 4B in Liu et al. (2006), Figure 4A in DeGiorgio et al. (2009), and Figure 3A in Balloux et al. (2009) (see Text S3). The BICs found with the quadratic models gave an origin area in just the south of Africa (Figures 9E and 10A). This adds to research (e.g., Henn, Cavalli-Sforza, & Feldman, 2012) in favour of the south being the start of the expansion. However, there were few populations at shorter distances (e.g., Figure 9D). Therefore, a lot of the non-linear trendline (0-2,000 km, i.e., about half of the range of distances in sub-Saharan Africa) covers just a few populations (e.g., Figure 9D). And so, there may be low confidence in the non-linear trendline being representative of how diversity changes with distance, and therefore low confidence in the origin area. Even if there is no initial increase, what may seem clearer is that the decline is not present across distances, and seems to be absent before approximately 2,000-3,000 km (e.g., Figure 9D). If the decline is present at shorter distances, then one may expect diversity to be far higher than it is before 2,000-3,000 km.

Why may there have been an increase at shorter distances (or a decline only being present after a few thousand km)? This could be explained by ideas in DeGiorgio et al. (2009) or Liu et al. (2006) (see Text S3), or somewhat build on both by focusing on admixture and population density. Indeed, wherever modern humans emerged from in Africa need not be the origin of their expansion across the continent (Henn, Bustamante et al., 2011). Indeed, it is conceivable that modern humans did not emerge from any one particular place, but across regions of Africa (e.g., Henn et al., 2018). An idea (amongst a number) is that there was expansion from a population into other areas within Africa in which modern humans were already located (with interpopulation gene flow occurring, therefore admixture) (Henn et al., 2018). And so, given that admixture is held to affect diversity (e.g., Skoglund & Mathieson, 2018), perhaps Figures 9D, 9E, and 10A point toward the initial part of the expansion having involved significant admixture between African populations, which resulted in diversity increasing at first. So, what would have been a loss of diversity in expansion (e.g., Ramachandran et al., 2005) would have been offset by admixture in the early part of expansion, such that the diversity either increased or stayed constant early on. Therefore, it could be suggested that the expansion was initially into more densely-populated areas in Africa (less than approx. 2,000-3,000 km from the peak point). The expansion scenario of bottlenecks *and* declining diversity could then be in effect in Africa (e.g., Prugnolle, Manica, & Balloux, 2005; Ramachandran et al., 2005) only from about 2,000-3,000 km onwards. That could be expansion to places which had not been inhabited by modern humans (e.g., Henn, Cavalli-Sforza, & Feldman, 2012). Or, maybe expansion could yet still have been to places which modern humans already were living in (e.g., Henn et al., 2018), but populated to a lesser extent compared to the earlier part of the expansion. Therefore, the influence of bottlenecks on diversity (e.g., Ramachandran et al., 2005) would become dominant only after around 2,000-3,000 km because there would be less admixing than earlier in the expansion. Alternatively, perhaps the apparent increase can be explained by variation in peak points between sets of populations (e.g., Figure 9A-9C); when the sub-Saharan African populations are grouped by regions used in Tishkoff et al. (2009), peak points seem to vary markedly (Figures 9B, 9C, and 10B). However, given expansions outside of the worldwide expansion (e.g., Hunley & Cabana, 2016), such as the Bantu expansion in the Holocene (Li et al., 2014), some interregional variation in peak points is surely to be expected. Moreover, few populations were used in southern and western regions (four and 10 populations respectively), so their peak points are certainly questionable.

### Autosomal SNP haplotype diversity: San

Compared to the other peak points (Ramachandran et al., 2005; Tishkoff et al., 2009; present research) which were used for Figure 6, the peak point for autosomal SNP haplotype heterozygosity (Figure 7D – larger circle) seems quite far north. The autosomal SNP haplotype heterozygosities of populations were sourced from a table in Balloux et al. (2009), who got these heterozygosities from Li et al. (2008). Li et al. (2008) calculated autosomal haplotype heterozygosities from chromosome 16 in HGDP-CEPH data. Citing research favouring ascertainment bias being lesser with SNP *haplotype* heterozygosity than *SNP* heterozygosity (i.e., Conrad et al., 2006), Li et al. (2008) used haplotype heterozygosity in regard to expansion. However, given preceding research (Conrad et al., 2006; Schlebusch et al., 2020), including research finding a decline of diversity as distance grows from Africa (e.g., Balloux et al., 2009; Prugnolle, Manica, & Balloux, 2005), Figure 3G of Balloux et al. (2009) and Table S3 of Balloux et al. could suggest that haplotype heterozygosity was unexpectedly low for the San population (but see López Herráez et al., 2009).^28^ Accordingly, when using an origin (10°S, 25°E) in the vicinity of the origin used by Balloux et al. (2009), San were found, in the present research, to have a low haplotype heterozygosity (with a standardised residual of −3.66).

San are the only one of the HGDP-CEPH populations (López Herráez et al., 2009) in southern Africa (Choudhury et al., 2021) of the HGDP-CEPH populations used in Balloux et al (2009) (and, therefore, in the present research) with respect to autosomal SNP haplotype heterozygosity.^29^ Therefore, bearing in mind how the origin area is calculated through BICs (Betti et al., 2009; Manica et al., 2007), and considering that populations in Africa seem to be given a considerable weighting when determining peak points (Figure 7; cf Van Tuyl & Pereltsvaig, 2012), low San heterozygosity may have led to a northward shift in the peak point and the origin area; without San, the origin area is indeed more southerly and the peak point moves slightly southward (Figure 1, Figure S7). As noted in Text S3, the unusuality of datapoints can vary depending on the origin used. When setting the origin as the peak point found without San, the San datapoint seems atypical (standardised residual of −3.32). And so, (also given Conrad et al., 2006) perhaps it could be argued that ascertainment bias remained to a sufficient level to affect the peak point and origin area for autosomal SNP haplotype heterozygosity.^30^

### Y-chromosomal microsatellite diversity: Within-Africa trend?

Y-chromosomal microsatellite heterozygosity appeared to be unrepresentative of expansion from Africa, instead pointing to worldwide expansion from Asia (e.g., Figure 5C and 5D). This was explored further. *Somewhere* being the origin of expansion is congruent with a *decline* of diversity as distance from that somewhere increases (e.g., Ramachandran et al., 2005). Research has taken place regarding whether a fall in diversity is indicated at the regional level (e.g., Hunley & Cabana, 2016; Hunley et al., 2016; Wang et al., 2007; see Text S2); a decline of autosomal microsatellite heterozygosity appears within Africa (in the present research), so, if Y-chromosomal microsatellite heterozygosity reflects expansion from Asia, one might expect a decline of Y-chromosomal diversity within Africa as distance from Asia increases. However, the negative correlation between distance from India and Y-chromosomal microsatellite heterozygosity might not be reflective of populations within Africa (but arise from populations outside of Africa). This can be said because when analysis was narrowed just to populations in Africa, then distance from India gave a positive correlation coefficient, *r*(5) = .70, unlike when populations outside of Africa were used, *r*(42) = −.61. Yet, when using distance from Africa (the location which had the lowest *sr* in Africa when Y-chromosomal diversity was adjusted for distance from India) instead of distance from India, the correlation coefficient was negative regarding populations inside of Africa, *r*(5) = −.90. And so, whilst it may appear that expansion from Asia is supported by Y-chromosomal microsatellite heterozygosity (e.g., Figure 5D), Y-chromosomal microsatellite heterozygosity within Africa *might* not be consistent with this.

Regarding the association of autosomal diversity and distance from Africa, Prugnolle, Manica, and Balloux (2005) grouped populations into categories, and compared gradients across the categories (gradient for Africans, gradient for Americans, etc.). However, as noted in Text S2, each category had few populations with the exception of one (Asians), which may have meant that there was a low likelihood of true differences between the gradients being apparent. As Prugnolle, Manica, and Balloux (2005) used HGDP-CEPH data (Lawson Handley et al., 2007; Ramachandran et al., 2005) (and from looking at Prugnolle, Manica, and Balloux, 2005, Balloux et al., 2009, and López Herráez et al., 2009), the seven populations used in Africa by Prugnolle, Manica, and Balloux (2005) were the same ones used in the present analysis. Therefore, a comparison of gradients was not undertaken in the present analysis (Africa compared to outside of Africa) because that analysis would be underpowered. Hence, the direction of correlation coefficients was focused on. Nevertheless, an analysis like in Prugnolle, Manica, and Balloux (2005) is what should be aimed for.

### Expansion origin centroid: Additional support?

Might findings with phonemic diversity (Atkinson, 2011; Van Tuyl & Pereltsvaig, 2012), nucleotide diversity (Luca et al., 2011), and linkage disequilibrium (Henn, Gignoux et al., 2011) bolster the origin centroid? The peak point concerning the negative relationship between phonemic diversity and geographical distance (Van Tuyl & Pereltsvaig, 2012) is in the vicinity of the overall peak point found with autosomal diversity (Figure 6). However, whilst Atkinson (2011) found that Africa exclusively holds the area that is within four BICs of the lowest BIC regarding phonemic diversity (phoneme inventory size), Africa was not included in such an area in Creanza et al. (2015). Indeed, Creanza et al. (2015) found the lowest BIC (or Akaike information criterion) was not even within Africa, but in Europe, with there being a negative association between phoneme inventory size and distance from Europe (see Creanza et al., 2015, for their thoughts on how the negative association came about). Generally, given previous research/discussion (Atkinson, 2011, 2012; Creanza et al., 2015; Van Tuyl & Pereltsvaig, 2012), it seems uncertain whether phonemic diversity is useful for indicating the origin within Africa of the worldwide expansion. Consequently, it is not clear if findings with phonemic diversity (Atkinson, 2011; Van Tuyl & Pereltsvaig, 2012) lend support to the origin centroid.

In Luca et al. (2011), correlation coefficients were calculated regarding nucleotide diversity and distance from locations in Africa. The (numerically) largest correlation coefficient regarding nucleotide diversity and geographical distance was when distances were from southern Africa (and this coefficient was negative) (Luca et al., 2011). Using only populations in Africa, in Henn, Gignoux et al. (2011), there was a positive correlation between linkage disequilibrium and distance from southwestern Africa. Furthermore, out of locations used as origins in Africa, it was found that the strongest positive correlation was within the southern-western area of Africa (Henn, Gignoux et al., 2011).^31^ However, it has been noted by López et al. (2015) that, in Schlebusch et al. (2012), linkage disequilibrium in South Africa was roughly as low as it is in places within Africa that are outside of South Africa. Given Schlebusch et al. (2012) and wording in López et al. (2015, p. 59), presumably López et al. (2015) were referring to southern Africa rather than the Republic of South Africa in particular.

### More caveats, and future research

#### Origin area estimation problem?

As discussed above, building on Van Tuyl and Pereltsvaig, (2012), when genetic/cranial data consist of populations across the globe, the peak point (within Africa) appears to disproportionately be due to populations within Africa. A disproportionate weighting may imply a problem when estimating the origin area. From examining a formula for calculating the BIC (Masson, 2011), the number of populations (no matter whether they are inside Africa or outside) would affect the size of the origin area – a larger number of populations would reduce the area in general.^32^ An impression from previous research is that, when estimating origin areas (Betti et al., 2009, 2013; Manica et al., 2007), it seems usual for few populations to be in Africa, with most populations being outside of Africa. In such research, the size of the estimated origin area would be affected by the number of populations outside of Africa (and within). It could be bothersome that, when using populations worldwide, the peak point appears to be determined disproportionately (not fully) by populations in Africa (Figure 7; cf. Van Tuyl & Pereltsvaig, 2012) whereas the size of the origin area would be notably contributed to by populations outside of the continent (and not largely by populations within Africa). Surely the inside Africa vs. outside Africa influence really *ought* to be similar for both the peak point and the size of the origin area. So, it could be argued that the size of the origin area may be smaller than it ought to be. This might not only be an issue regarding the BIC. Origin areas estimated through the Akaike information criterion (AIC) (e.g., Betti et al., 2013) may also be affected. Consequently, estimated origin areas in prior research (e.g., Betti et al., 2013; Manica et al., 2007), and several in the present research, could be smaller than they should be. Nevertheless, this possible problem with origin areas was avoided in the analyses which used populations in sub-Saharan Africa alone (Figure 8).

For worldwide samples, a remedy to the possible estimation problem can perhaps be found in Tishkoff et al. (2009). Through a bootstrapping method, they represented on a map the frequency with which a location was the strongest (numerically) for the relationship between geographical distance and autosomal microsatellite diversity in repeat lengths (Tishkoff et al., 2009). The resulting area (Tishkoff et al., 2009) would likely not be subject to the underestimation which may affect BIC-based and AIC-based methods.

#### Effect of African populations

It is hoped that future research will go further than either previous research (Van Tuyl & Pereltsvaig, 2012) or the present research when exploring if African populations are the main influencers of peak points. In the present research, there was a visual comparison between peak points found with populations worldwide and in Africa only. It may seem like populations in Africa are the main (yet not sole) determinants of peak points for populations worldwide (Figure 7; cf. Van Tuyl & Pereltsvaig, 2012).

Perhaps a further approach could involve analysing patterns in correlation coefficients regarding the locations (in Africa) which were made use of like they are origins (Figure 11). If African populations are great determiners of peak points, one may expect a pattern between correlation coefficients found for African populations and coefficients for populations worldwide, with the pattern resembling a line. Such a pattern looks to be reasonably apparent for cranial shape diversity and cranial size dimorphism, but to a lesser extent for genetic diversities, in particular mitochondrial diversity for which the distribution of correlation coefficients seems quite different (Figure 11). This adds weight to Figure 7, for instance regarding the discrepancy between the mitochondrial diversity peak points for African populations and populations worldwide. Therefore, whilst exploring the influence of populations on peak points, future research could see if this influence is constant between variables.

**Figure 11.**
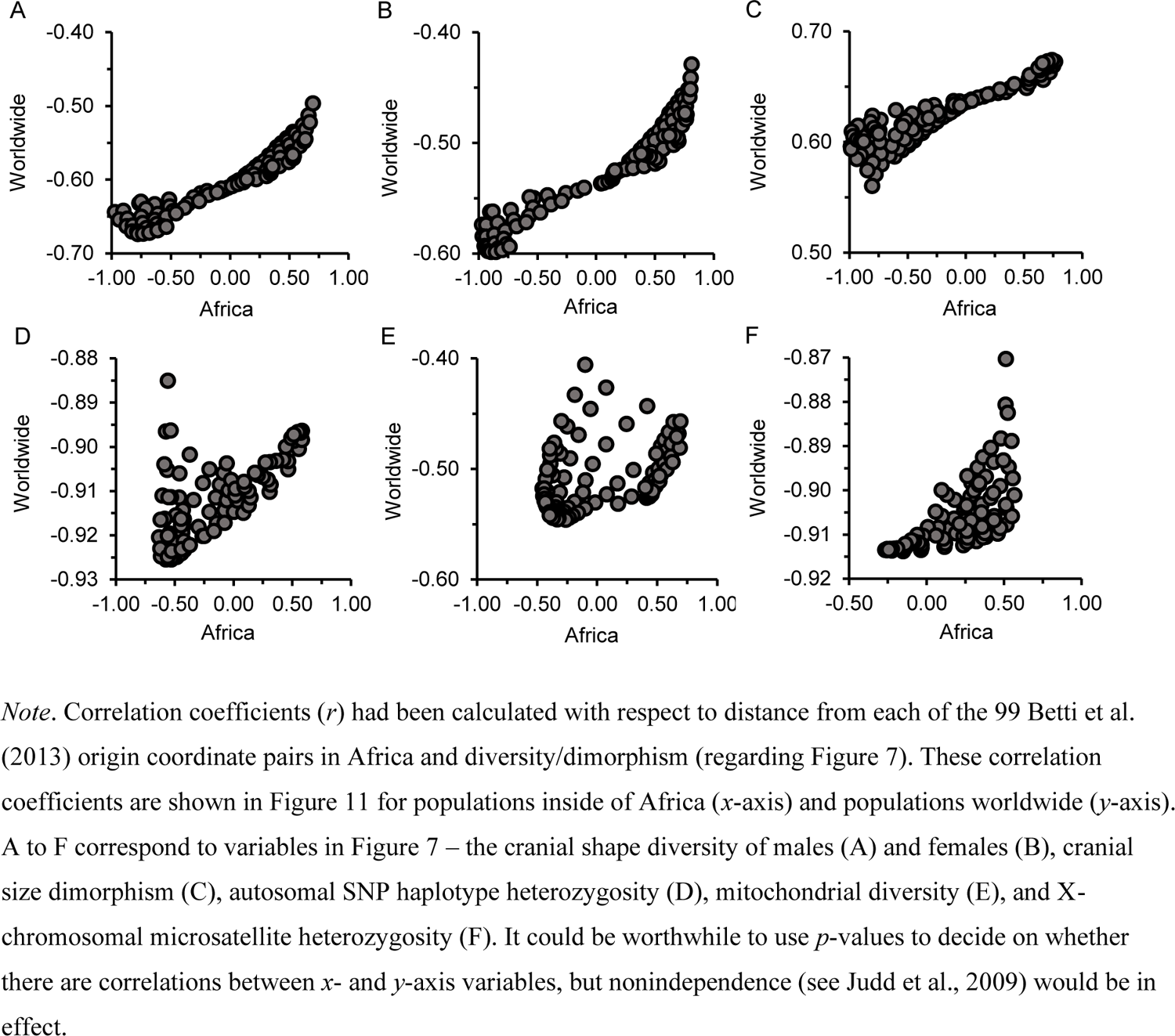
Correlation Coefficients: Populations in Africa, and Population Worldwide

#### Defining which variables signal the expansion

In the present research, for a variable to be noted as an indicator of the expansion from Africa and be included when calculating the origin centroid, certain criteria were applied; the variable had to correlate with distance from Africa, and the variable had to signal an origin area that is only within Africa. These criteria may be biased in favour of variables that have peak points which are more southerly/westerly within Africa. Why may there be a bias? The answer to this question is in regard to the size of the origin area.

An origin area can encompass quite a lot of continental Africa (e.g., Figure 2). Therefore, given support for a northeastern exit from Africa in the worldwide expansion (Tishkoff et al., 2009), a variable with a peak point (in Africa) closer to this exit is perhaps more likely to have an origin area which reaches outside of Africa. Consequently, with respect to peak points in Africa, a variable that has its peak point nearer to the exit may have a higher likelihood of being discarded as an indicator of the expansion. And so, a bias may have occurred when applying criteria used in the present research for defining an indicator of the expansion. Therefore, the origin centroid (Figure 6) could have been biased against peak points which are nearer to the exit. This bias could promote finding a southerly/westerly centroid in Africa. Indeed, the centroid is in southwestern Africa (Figure 6).

More relaxed criteria might reduce the possible bias. It seems that, in Betti et al. (2013), it was acceptable for diversity in hip bone shape to be considered as being indicative of expansion from Africa despite Betti et al. having found that hip bone shape diversity has an origin area which is not entirely situated in Africa. Indeed, hip bone shape diversity falls with the increase in geographical distance from Africa, it has a peak point in Africa, and whilst it leads to an origin area not fully within Africa (Betti et al., 2013), Figure 4 of Betti et al. (2013) shows that the area is mostly in Africa; the relaxed criteria could be i) an association with distance, ii) a peak point in Africa, and iii) an origin area at least largely in Africa. With hip bone shape diversity, the peak point is in southern Africa for males and for females (Betti et al., 2013). Hence, even with a possible bias in calculating the origin centroid, this possible bias might not necessarily have greatly affected which region of Africa the centroid is located in.

#### Mitochondrial diversity and climate

Whereas previous research has found an origin area and peak point using cranial form diversity after adjusting this type of cranial diversity for climate (Manica et al., 2007), research does not support cranial form diversity (overall) being associated with climate (Betti et al., 2009) (although climate is associated with the diversity of some cranial dimensions, Manica et al., 2007). There could be good reason to adjust mitochondrial diversity for climate. Mitochondrial diversity is associated with minimum temperature (Balloux et al., 2009). Furthermore, mitochondrial diversity (with distance from Africa being adjusted for) increases with minimum temperature (Balloux et al., 2009). Hence, the origin area (Figure 4) and peak point found with mitochondrial diversity (Figure 6) could perhaps be refined by adjusting mitochondrial diversity for minimum temperature.

However, the distribution of population locations in Africa may need to be kept in mind. The peak point found with (unadjusted) mitochondrial diversity is in southern Africa when using populations from across the globe (Figure 6). However, when analysing mitochondrial diversity in the present research, going by Balloux et al. (2009), no populations in southern Africa were featured. With this absence of southern African representation, there is reason to be hesitant about both the origin area and peak point for mitochondrial diversity.

#### Admixture and diversity

Previous research indicates that SNP heterozygosity in Africans decreases as Eurasian ancestry greatens (Haber et al., 2016). It could be worthwhile considering if this (to some extent) could be explained by bottlenecks within Africa in the worldwide expansion given that the present research suggests i) that the worldwide expansion began in southern Africa and ii) that the signal of the expansion is present in Africa. Bearing in mind research regarding Native Americans (Hunley & Healy, 2011), it would not be surprising if support for the association between Eurasian ancestry and genetic diversity in Africa (Haber et al., 2016) would still be indicated despite the expansion signal in Africa. In that previous research with Native Americans, there was a negative correlation between gene identity (of autosomal microsatellites) and European ancestry, and a negative correlation was still present when adjusting for the distance between Beringia and populations (Hunley & Healy, 2011).

#### Type of origin

The geographical area which is indicated to be the one from which modern humans expanded (i.e., expansion origin) can be interpreted as being the suggested area for where modern humans *originated* (i.e., where they first lived – modern human origin) (Manica et al., 2007); the origin centroid could be described as signaling the geographical location at which modern humans did originate. Regarding populations in Africa, it is possible that their ancestors were outside of southern Africa (but still within Africa) prior to going to the south and expanding (Henn, Bustamante et al., 2011). Therefore, origin areas for the worldwide expansion in the present analyses (e.g., Figure 1A; Figure 2A) need not imply that modern humans originated within those specific areas, and the expansion origin centroid (Figure 6) is not necessarily suggestive of modern humans having originated in southern Africa. Furthermore, the present research is not intended to suggest that all the ancestry of modern humans descends from the geographical origin of the worldwide expansion (for instance, see the literature on Denisovan/Neanderthal ancestry, e.g., Green et al., 2010; Yuan et al., 2021).

#### Within sub-Saharan Africa

In the present research, the autosomal microsatellite heterozygosity of sub-Saharan Africans was focussed on in some analyses. This was done in order to reduce a conceivable influence of non-African ancestry in Africa when studying the expansion in order to infer whether the worldwide expansion signal is present within Africa. In Schlebusch et al. (2012), genetic variables in sub-Saharan African populations (e.g., haplotype diversity) were absent of any distinct shift over geography. Whilst Schlebusch et al. (2012) did not examine whether autosomal microsatellite heterozygosity (of sub-Saharan Africans) changes with distance from southern Africa, the contrast between their results and the present research should call for more exploration of whether diversity in sub-Saharan Africa expresses the worldwide expansion.

And so, it is hoped that the approach taken with autosomal diversity in the present research (i.e., using sub-Saharan Africans only) will be applied to additional variables in future research. For instance, cranial form/shape diversity indicates the expansion (Betti et al., 2009; Manica et al., 2007; von Cramon-Taubadel & Lycett, 2008; Figure 2). Based on research with cranial diversity (Betti et al., 2009; Manica et al., 2007; von Cramon-Taubadel & Lycett, 2008; present research), the greatest number of sub-Saharan African populations in an analysis (when studying cranial diversity and the expansion) was in Manica et al. (2007) and Betti et al. (2009). Their analyses featured only 12 sub-Saharan African populations amongst males (and only five of such populations for females) (Betti et al., 2009; Manica et al., 2007) – this seems few compared to the 106 used in the present study for autosomal diversity. The cranial diversity used in Manica et al. (2007) and Betti et al. (2009) was cranial form diversity; Betti et al. (2009) obtained data from Manica et al. (2007). If one uses, from Betti et al. (2009), the cranial form diversity of male sub-Saharan African populations and distances from Africa, a correlation coefficient of *r* = −.36 is found regarding these two variables. In Betti et al. (2009), South African populations had a distance (from sub-Saharan Africa) of under 300 km – the distances in Betti et al. appear to have been from southern Africa. From reading Betti et al. (2009), it is not clear whether those distances were from the peak point for sub-Saharan African populations; if the distances were not from the peak point, perhaps a numerically lower coefficient than −.36 would be found using distances from the peak point.^33^ Nevertheless, the coefficient seems comparable to the correlation coefficient observed in the present research regarding the autosomal diversity of sub-Saharan Africans (in parametric analysis with no adjustment for non-African ancestry). Cranial form diversity may be less effective at indicating the origin of expansion than cranial shape diversity (Figure 2). Therefore, it could be particularly useful to employ cranial shape diversity when exploring if the diversity of sub-Saharan Africans indicates the expansion – providing one has a sufficient number of populations.

To estimate where the expansion originally came from, ideally, it could be seen where peak points cluster and where a centroid is located (e.g., like in Figure 6), but solely using sub-Saharan African populations. For a variable known to indicate the expansion (e.g., von Cramon-Taubadel & Lycett, 2008; Figure 2), if few sub-Saharan African populations are featured, then the variable could potentially be included when evaluating the locations of peak points and the centroid – looking for a general suggestion of an origin across variables may alleviate the problem of having few sub-Saharan African populations in a variable.

## Conclusion

The present analyses explored which cranial and genetic variables may signal the worldwide expansion of modern humans, and where in Africa this expansion commenced from. Support was found for the signal being present in various types of diversity (e.g., X-chromosomal microsatellites, cranial shape) and cranial size dimorphism. Nonetheless, despite Y-chromosomal microsatellite heterozygosity being related to geographical distance from Africa (Balloux et al., 2009), that type of diversity was not indicated to signal the expansion. Yet that diversity was from six Y-chromosomal microsatellites (Balloux et al., 2009), i.e., not many (given Seielstad et al., 1999), so it really is not clear if the signal is present in the heterozygosity of Y-chromosomal microsatellites. As for where in Africa the expansion originated, the location of a geographical centroid (determined through a number of expansion indicators) and locations of peak points (using populations worldwide) adds weight to southern Africa being the origin (Figure 6). In the context of research with phonemic diversity (Van Tuyl & Pereltsvaig, 2012), the locations of these peak points and this centroid may greatly arise from biological diversity/dimorphism within Africa (Figure 7). This points towards the signal of the expansion being found amongst populations in the continent. Indeed, analyses with autosomal microsatellite diversity in sub-Saharan African populations suggested the presence of the expansion signal in Africa, with the expansion seeming more likely to have originated in southern Africa than in other regions of Africa. Even with the support for a non-monotonic relationship between autosomal microsatellite heterozygosity and distance within Africa, a southern African origin is indicated. Nevertheless, a number of caveats (as described above) do remain. It is far from a certainty that southern Africa is the region from which the worldwide expansion originated.

### Box 1

#### Glossary

- **Expansion origin centroid**: estimated from different variables, this is an estimate of where the expansion first spread from (i.e., where it originated) – for an example, see Figure 6.
- **Geographical area of best fit**: a geographical area made totally of locations from which distances have the greatest of associations with a variable – for examples, see Betti et al. (2009) or Figure 1A.
- **Origin area (area of origin)**: a geographical area of locations amongst which the origin of the expansion is estimated to be – for examples, see Manica et al. (2007) or Figure 2A.
- **Peak point**: the *best* (singular) location in Africa for being the origin of the expansion – for examples, see Figure 7.

## Footnotes

^1^ Indeed, in accordance with expansion from Africa are declines in, for example, autosomal SNP haplotype heterozygosity (Li et al., 2008), autosomal microsatellite repeat length diversity (Tishkoff et al., 2009), cranial form diversity (Manica et al., 2007), cranial shape diversity (von Cramon-Taubadel & Lycett, 2008), pelvic shape diversity (Betti et al., 2012), and hip bone shape diversity (Betti et al., 2013).

^2^ Indeed, depending on which location is utilised as the origin, correlation coefficients on the association between diversity and geographical distance can vary between negative and positive (e.g., Hunley et al., 2012; Ramachandran et al., 2005). An example of this can be found with autosomal microsatellite diversity – the correlation coefficient is negative when using Africa as an origin, whilst the coefficient is positive when South America is used (Ramachandran et al., 2005).

^3^ Autosomal microsatellite heterozygosity, for example, achieves points i (Balloux et al., 2009; Prugnolle, Manica, & Balloux, 2005) and ii (Manica et al., 2007), and it has been used to explore where the expansion had its beginning (Manica et al., 2007; Ramachandran et al., 2005).

^4^ With female cranial diversity in form (39 populations [*n* = 1,579]), the origin area is not in Africa alone (Betti et al., 2009). Indeed, for females, the origin area is wider than it is for males, however, the sample size for females was not large in that study (Betti et al., 2009). Regarding hip bone shape diversity, the origin area is not exclusive to Africa for males (27 populations [*n* = 891]) or females (20 populations [*n* = 552]) (Betti et al., 2013).

^5^ Nevertheless, form encompasses size and shape aspects (Betti et al., 2010), and previous research observed that cranial size diversity, adjusted for absolute latitude, and distance from Africa have no correlation (Cenac, 2022), so it would be sensible to assume that the origin area found with cranial shape diversity would be in Africa alone for males, and perhaps for females too.

^6^ Given Balloux et al. (2009) and some research (i.e., Prugnolle, Manica, & Balloux, 2005; Prugnolle, Manica, Charpentier, et al., 2005) cited by them, it can be interpreted that the declines in X-chromosomal, Y-chromosomal, and mitochondrial diversity were shown by Balloux et al. to be consistent with expansion from Africa.

^7^ Figure 1 of Henn, Cavalli-Sforza, and Feldman (2012) represents expansion of modern humans from inside Africa to Eurasia, and from Eurasia to Oceania and the Americas.

^8^ Simulations in Ray et al. (2005) were regarding data which Rosenberg et al. (2002) utilised – Rosenberg et al. used data from autosomal microsatellites.

^9^ After referring to previous research (specifically Prugnolle, Manica, & Balloux, 2005, and Ramachandran et al., 2005) on autosomal microsatellite heterozygosity falling as distance from eastern Africa increases, Deshpande et al. (2009) appear to have noted that the decline is strongest from East Africa. However, Prugnolle, Manica, and Balloux (2005) did not describe examining whether a decline in autosomal microsatellite diversity is greater when distance is from East Africa rather than anywhere else. Additionally, Ramachandran et al. (2005) did not find that the peak decline occurs when using distance from East Africa. Moreover, Manica et al. (2007) estimated an origin area using autosomal microsatellite heterozygosity; that origin area did include East Africa, but also other regions of Africa (Manica et al., 2007).

^10^ Additionally, Colonna et al. (2011) can be regarded as referring/alluding to origin estimation in Africa being affected by post-expansion admixture and population movement in Africa.

^11^ Research supports there being a bottleneck regarding exit from Africa (Hellenthal et al., 2008).

^12^ *Z*-scores of residuals (i.e., standardised residuals) can be used when assessing if datapoints are atypical – for instance, datapoints exceeding 3.29 in terms of their absolute value may be atypical (e.g., Field, 2013), and that value for absolute *z*-scored residuals was indeed used for identifying atypical datapoints in parametric analyses within the present research.

^13^ The names given for populations in the Howells data (Howells, 1989, 1996) are used in the present research.

^14^ Research/manuals (Excoffier, 2004; Excoffier & Lischer, 2015; Marjoram & Donnelly, 1994; Schneider et al., 1997; Slatkin & Hudson, 1991) were consulted by the author of the current manuscript for clarification regarding the mitochondrial diversity.

^15^ Regarding the HGDP-CEPH and mitochondrial genetic data, the calculation of geographical distances in the present research involved using population coordinates obtained from Balloux et al. (2009) alongside maps in Bartholomew illustrated reference atlas of the world (1985) and López Herráez et al. (2009) in order to be clear on the continent of each population – for mitochondrial data, continents of populations largely matched continental assignments in Balloux et al. (2009). Regarding HGDP-CEPH populations, continents had been stated in Tishkoff et al. (2009), Pemberton et al. (2013), and Rosenberg et al. (2002), but were not a factor here. Continents with respect to the Howells cranial data were regarded as in Cenac (2022).

^16^ Von Cramon-Taubadel and Lycett (2008) used three origins in Africa, and two of those origins feature amongst the 99 African origins in Betti et al. (2013). And so, two of the African origin coordinate pairs used in von Cramon-Taubadel and Lycett (2008) were also used in the present research (due to them being part of the 99) when constructing origin areas.

^17^ Continental groupings of origins in graphs were as in Cenac (2022).

^18^ Extents of African and non-African ancestry are shown in Figure 3 of Tishkoff et al. (2009).

^19^ The intention was to list, in column two, only research which included a *p*-value (with respect to distance from Africa having an association with a variable). The author of the present research thought such a *p*-value was given in Tishkoff et al. (2009), but, upon looking through Tishkoff et al. (2009) again, may have been mistaken. So, criteria regarding column two was changed – research concerning a fall/rise in a variable as distance from Africa ascends did not have to include a *p*-value.

^20^ In the present analyses, distance from Africa (Botswana) was found to have a negative correlation with cranial shape diversity amongst males, *r*(26) = −.67, *p* = .001, spatial *DW* = 2.39. Because von Cramon-Taubadel and Lycett (2008) had already found a correlation coefficient and *p*-value regarding distance from Botswana and cranial shape diversity using the male crania (28 populations in the Howells dataset), it was not necessary for this correlation test to have been run in the current analyses. However, von Cramon-Taubadel and Lycett (2008) did not state whether they examined residuals for positive spatial autocorrelation.

^21^ Given preceding research using linkage disequilibrium (regarding autosomal SNPs) (Henn, Gignoux et al., 2011; Jakobsson et al., 2008), it is clear that cranial size dimorphism is by no means unique in having a positive association with distance from Africa. Indeed, amongst populations worldwide, linkage disequilibrium grows with distance from Africa (Jakobsson et al., 2008). Moreover, inside of Africa, linkage disequilibrium does increase with distance, and an origin area in the southwestern region has been found using linkage disequilibrium (Henn, Gignoux et al., 2011).

^22^ To clarify, Tishkoff et al. (2009) did find their peak point (autosomal microsatellite repeat length diversity) to be around the Angola/Namibia border – in the vicinity of the seaboard.

^23^ Tishkoff et al. (2009) used worldwide populations, including 121 in Africa. The peak point in their study for the relationship between geographical distance and diversity in autosomal microsatellite repeat lengths (Tishkoff et al., 2009) is close to the origin centroid of Figure 6 (in the present research, that peak point was used to calculate the centroid for autosomal diversity, which was used to arrive at the origin centroid).

^24^ Research/discussion has occurred regarding genetic diversity in sub-Saharan Africa and geography (e.g., Bergström et al., 2021; Schlebusch et al., 2012). Present analyses with autosomal microsatellite heterozygosity in Africa only used populations in sub-Saharan Africa. This is because of research in line with i) migration back into the continent of Africa (Henn, Botigué et al., 2012; Hodgson et al., 2014), ii) the presence of non-African ancestry in the continent (Tishkoff et al., 2009), and iii) for Africans with admixture, heterozygosity falling as Eurasian ancestry increases (Haber et al., 2016). Therefore, solely populations in sub-Saharan Africa were used in order to lower any impact on results of migration back into Africa. That said, non-African ancestry in sub-Saharan Africa (e.g., Tishkoff et al., 2009) is acknowledged.

^25^ Ancestry has been adjusted for (Hunley & Cabana, 2016; Hunley et al., 2016) previously. For instance, research on expansion has adjusted distance from Africa and the autosomal diversity of Native Americans for European ancestry (Hunley & Cabana, 2016).

^26^ Previous research explored correlations between gene identity and European ancestry amongst Native Americans when adjusting both variables for how far from Beringia populations are, and when not adjusting variables for distance (with gene identity being the number one minus gene diversity) (Hunley & Healy, 2011); in the present research, also given the research aligning with a relationship between Eurasian ancestry and the heterozygosity of Africans (Haber et al., 2016), the Tishkoff et al. (2009) data was utilised to find if non-African ancestry correlates with the autosomal heterozygosity of populations in sub-Saharan Africa. The autosomal heterozygosity was adjusted for distance from the peak point in some correlation tests. Interestingly, there was no correlation between the non-African ancestry and heterozygosity, whether heterozygosity was, *sr*(103) = .13, *p* = .72, spatial *DW* = 2.00 [*sr*_s_(103) = .17, *p* = .44] or was not adjusted for distance from the peak point, *r*(104) = .09, *p* = .97, spatial *DW* = 1.89 [*r*_s_(104) = .10, *p* = .97]. There appears to have been little variability in the non-African ancestry (Figure S6), and this lack of variation could explain the absence of correlations.

^27^ Using populations across the world, a quadratic trend (increase than decrease) has been found between distance from Africa and what is arguably an indicator regarding economic development (Ashraf & Galor, 2013) (but see Rosenberg & Kang, 2015 and d’Alpoim Guedes et al., 2013).

^28^ Indeed, in Li et al. (2008), it can be seen that autosomal SNP *haplotype* heterozygosity in HGDP-CEPH data is numerically lower for San than for populations of Europe or the Middle East (also, in Li et al., 2008, with respect to *SNP* heterozygosity, this diversity was described as not being as great in San as in those other populations); as for autosomal and X-chromosomal microsatellite heterozygosities, in Balloux et al. (2009), San were numerically greater than those populations.

^29^ Using the classification of southern Africa in Choudhury et al. (2021), it is clear from Shi et al. (2010) that San are not the sole HGDP-CEPH population in southern Africa.

^30^ San haplotype heterozygosity seems relatively lower amongst HGDP-CEPH populations in Li et al. (2008) (autosomal) than in Conrad et al. (2006) (regarding results shown at the largest window size in their Figure 3) who used autosomal and X-chromosomal data.

^31^ Moreover, Henn, Gignoux et al. (2011) used genetic distances between each of 13 African genetic clusters and Italians, and geographical distances. In Henn, Gignoux et al. (2011), the south as the origin for dispersal in Africa seemed to be supported based on correlation coefficients when various locations were rolled out as geographical origins. Yet, the regression regarding geographical distances (from the particular origin which granted the most negative of the correlation coefficients) and genetic distances resulted in a *p*-value which was over .05 (Henn, Gignoux et al., 2011). However, just 13 datapoints were used in the regression analysis (Henn, Gignoux et al., 2011).

^32^ In a previous study, BIC was used for finding the origin area through cranial form diversity (adjusted for climate) (Manica et al., 2007). Regarding the crania of males (105 populations; 4,666 crania) and females (39 populations; 1,579 crania), amongst females, the dataset was too small to sift between possible origins (Manica et al., 2007) which suggests that the sample size influences the size of the origin area estimated through the BIC. ^33^ In Betti et al. (2009), analyses (which included populations globally) used locations as origins which swept across Africa. Therefore, Betti et al. (2009) would have used the peak point for sub-Saharan African populations at some point.

## Supporting information

Supplementary texts, table, figures, and references

