## Supplementary texts, table, figures, and references for "Modern human expansion from (southern?) Africa: Origin, non-African ancestry, and linearity"

**Supplementary materials**

**Text S1: Cranial size dimorphism and absolute latitude**

Some research on cranial form diversity and geographical distance has adjusted diversity for climate (Manica et al., 2007). A subsequent study suggested that cranial form diversity (mean diversity of cranial dimensions), which does have an association with distance from Africa, is not tied to climate (Betti et al., 2009); when analysing if cranial size dimorphism shares a correlation with distance from Africa, cranial size dimorphism has been controlled for absolute latitude (Cenac, 2022) without finding if there is any utility in making this adjustment, or establishing whether cranial size dimorphism is related to absolute latitude. In the present research, it was explored whether controlling cranial size dimorphism for absolute latitude is worthwhile, and if cranial size dimorphism is correlated with absolute latitude.

The zeroEQpart package (Richard, 2018), in R, was used to examine if two (extents of) correlations are of the same strength. The two correlations of interest were the correlations between distance from Africa and i) cranial size dimorphism (*zero-order correlation*), ii) cranial size dimorphism, with this dimorphism being adjusted for absolute latitude (*semi-partial correlation*). A default of 1,000 bootstrapped samples (Richard, 2018) featured in the analysis with zeroEQpart. The ppcor package (Kim, 2015) was used in R to test if cranial size dimorphism is correlated with absolute latitude when controlling cranial size dimorphism for distance from Africa. Given that research suggests a possible connection between (adjusted) cranial size dimorphism and expansion from Africa (Cenac, 2022), adjusting cranial size dimorphism for distance from Africa should sufficiently explain variation in cranial size dimorphism such that any ability of absolute latitude to relate to cranial size dimorphism may be made more apparent by this adjustment (compared to seeing if there is a zero-order correlation between absolute latitude and cranial size dimorphism).

***Data***

Cranial size dimorphism of 26 populations (adjusted for absolute latitude, and unadjusted) was calculated beforehand using the Howells (1973, 1989, 1995, 1996) data (that data had been gotten from a link on http://web.utk.edu/~auerbach/HOWL.htm) (Cenac, 2022), and this adjusted/unadjusted dimorphism was used in the Text S1 analyses. Absolute latitudes had been used beforehand via the latitudes presented in von Cramon-Taubadel and Lycett (2008)[1] (Cenac, 2022), and these absolute latitudes were utilised in the current analyses, as were geographical distances calculated in Cenac (2022) and (as described in *Method*) the present research.

***Correlation comparison***

Regarding the correlation test between adjusted (for absolute latitude) cranial size dimorphism and distance from Botswana, it is uncertain if residuals were positively spatially autocorrelated,

[1] Regarding the latitudes in von Cramon-Taubadel and Lycett (2008), negative latitudes had their sign switched to positive and the positive latitudes remained the same (Cenac, 2022).

however, positive spatial autocorrelation was absent when Ainu and North Japan were not featured (Cenac, 2022). Using ppcor (Kim, 2015), without the two populations, adjusted cranial size dimorphism was found to have a positive (semi-partial) correlation with distance from Botswana, $sr(21) = .66$, $p = .004$.[2] Through zeroEQpart, it was found that this correlation was just as strong as the zero-order correlation between distance from Botswana and cranial size dimorphism, $p = 1.00$. Hence, it would not seem useful to adjust cranial size dimorphism for absolute latitude when finding if this dimorphism and distance from Africa are correlated.[3]

***Bayesian information criterion***

The comparison analysis above did not use the peak point (either for adjusted or unadjusted cranial size dimorphism). Just because there was no difference in correlations (unadjusted vs. adjusted) when using distance from Botswana does not mean that there would be no difference if a peak point was used. Furthermore, adjusting for cranial size dimorphism could potentially change the distribution of BICs compared to when unadjusted cranial size dimorphism is used – peak points may not be the same.

Therefore, using the four-BIC criterion (e.g., Manica et al., 2007), it was inferred whether any of the BICs within Africa found with adjusted cranial size dimorphism were markedly lower than the BIC found with the peak point regarding unadjusted cranial size dimorphism (i.e., if any fits were better with adjusted dimorphism than with unadjusted dimorphism). Concerning unadjusted cranial size dimorphism, the BIC found using the peak point (which had been calculated in R – also see *Method*) was recalculated using a formula in Masson (2011) (that formula had been used regarding adjusted dimorphism, and BICs would appear to be calculated differently in R compared to the formula).

The peak point found with adjusted cranial size dimorphism was the same as the one found with unadjusted dimorphism. BICs did not indicate that distance from the peak point was a better fit to adjusted dimorphism than to unadjusted dimorphism. Therefore, adjusting cranial size dimorphism for absolute latitude would indeed appear to be unnecessary.

---

[2] Previous research had used the Spearman correlation route to see if there is a correlation (Cenac, 2022).

[3] Furthermore, absolute latitude did not correlate (zero-order) with cranial size dimorphism, $r(24) = .16$, $p = 1.00$, spatial $DW = 1.50$. Regarding a correlation test between absolute latitude and cranial size dimorphism (the latter controlled for distance from Botswana), it was uncertain if positive spatial autocorrelation was present when 26 populations were used, spatial $DW = 1.30$. A spreadsheet, which includes a graph, from Chen (2016), was inputted with data in the current research – the resulting graph indicated that this uncertainty resulted from Ainu and North Japan populations. Using the other 24 populations, positive spatial autocorrelation was not present, spatial $DW = 1.90$, and there was no correlation between cranial size dimorphism (which had been adjusted) and absolute latitude, $sr(21) = .06$, $p = 1.00$. Therefore, distance from Africa, rather than absolute latitude, had a relationship with cranial size dimorphism.

**Text S2: Expansion signals within regions?**

The fall of autosomal microsatellite heterozygosity (with the rising of distance from Africa) is in line with there being a serial founder effect (Ramachandran et al., 2005). Under a serial founder effect, more bottlenecks have been experienced with distance from the origin of the expansion (e.g., Sugden & Ramachandran, 2016). Atkinson (2011) applied the serial founder effect concept to language. Atkinson (2011) found that phonemic diversity has a decline with the furthering of distance from Africa, and (via phonemic diversity) found an origin area for language that was entirely in Africa (cf. Creanza et al., 2015). At a regional level, when it comes to phonemic diversity and distance from western Africa, the gradient is in the negative direction within Africa, yet not within regions that are outside of Africa (northern Asia, North America, South America) (Van Tuyl & Pereltsvaig, 2012); it has been questioned why a serial founder effect should be thought to underlie the gradient within Africa (Van Tuyl & Pereltsvaig, 2012). Therefore, if a decline in autosomal diversity is only present within Africa,[4] it may appear doubtful that the decline within Africa is a product of the expansion.

***Autosomal microsatellites***

The remainder of Text S2 focusses on i) research examining if population categories differ in the gradient regarding distance from Africa and autosomal microsatellite diversity (Prugnolle et al., 2005), and ii) studies seeing if there is, at a regional level, a relationship between geographical distance and autosomal microsatellite diversity (Hunley & Cabana, 2016; Hunley et al., 2016; Wang et al., 2007) or autosomal microsatellite gene identity (Hunley & Healy, 2011). The previous research (using autosomal microsatellite data) explored mean expected heterozygosity (Prugnolle et al., 2005; Wang et al., 2007), gene diversity (Hunley & Cabana, 2016; Hunley et al., 2016), or gene identity (Hunley & Healy, 2011) – the mean expected heterozygosity *is* gene diversity (Yang et al., 2010), and gene identity is gene diversity subtracted from the number one (Hunley & Healy, 2011).

*Differences between regions?*

Prugnolle et al. (2005) observed that autosomal microsatellite heterozygosity falls as distance from eastern Africa increases. The decline is indicated to be a consistent decline (Prugnolle et al., 2005). Indeed, this decline was the same across categories of populations (six of which were used, including African) in terms of gradients (and intercepts) (Prugnolle et al., 2005). Therefore, one may expect there to be negative correlations between distance from Africa and autosomal diversity inside of different continents. However, certain analyses in Prugnolle et al. may have been underpowered, i.e., too low a chance of finding actual differences. Prugnolle et al. (2005) used 51 populations located across the world. From Figures 1 and 2 in Prugnolle et al. (2005), it can be observed that each category featured few populations aside from the Asian category. For instance, the African population

[4] Or if a decline outside Africa can be attributed to admixture (Hunley & Cabana, 2016).

category had just seven populations (Prugnolle et al., 2005). So, the chance of finding differences in gradients and in intercepts (if there are differences) possibly was low.

*Africa*

Within Africa, Hunley and Cabana (2016) did not find relationships between distance and autosomal microsatellite gene diversity (three population categories in Africa were used separately). However, as explained in the main text, geographical distance was from eastern Africa rather than southern Africa – results in the present research support the origin being in the south.

*Americas*

Regarding the Americas, in Wang et al. (2007), support was found for autosomal microsatellite heterozygosity declining as the distance between populations and the Bering Strait increases amongst Native Americans, and this was not indicated to arise from European admixture. However, whilst Hunley and Cabana (2016) found that distance (Beringia to populations) is associated with autosomal microsatellite gene diversity amongst Native Americans, the *p*-value (numerically) exceeded .05 in their study when both distance and diversity were adjusted for European ancestry. Moreover, Hunley and Healy (2011) observed that autosomal microsatellite gene identity increases with distance from Beringia, yet not when adjusting for European ancestry. Nevertheless, Hunley et al. (2016) found the fall in autosomal microsatellite gene diversity of Native Americans (as distance increases from Beringia) remained after adjusting for non-Native-American (African, East Asian, European) ancestry. Therefore, given this research concerning the Americas (Hunley & Cabana, 2016; Hunley et al., 2016; Hunley & Healy, 2011; Wang et al., 2007), it seems questionable exactly what the relationship between distance and autosomal microsatellite diversity within the Americas (e.g., Hunley & Cabana, 2016) is likely to reflect – the worldwide expansion signal or admixture.

*Asia, Europe, and Oceania: Range of distances*

Hunley and Cabana (2016) found no correlation between distance from Africa and autosomal microsatellite gene diversity within Europe, or within Oceania, or within each of three regions in Asia. However, given a comment in previous research concerning phonemic diversity (Atkinson, 2012), could the range of geographical distances be a factor in whether correlations (regarding genetic diversity) are found at a regional level? In Hunley and Cabana (2016), going by Figure 1 in their study, the distances from Africa were between about 15,000 km to over 25,000 km for populations in the Americas region (so a range of around 10,000 km). That same Figure 1 in Hunley and Cabana (2016) suggests that the ranges regarding other regions used in Hunley and Cabana (e.g., Europe, Oceania) were notably less than the circa. 10,000 km range of the Americas. Regarding the decline of phonemic diversity with the increase of distance from Africa, statistical power is considerably lowered when searching for patterns inside continents (Atkinson, 2012). So, the ability to find a decline in autosomal diversity should be lowered when there is a smaller range of distances from Africa. Asia, in Hunley and Cabana (2016), could be used as an example of this. In Hunley and

Cabana (2016), Asia was divided into three regions. Figure 1 of Hunley and Cabana (2016) indicates that each of those three regions had a geographical range less than the Americas. Based on Figure 1 in Hunley and Cabana (2016), it seems possible that there would be a correlation between distance from Africa and diversity within Asia if Asia is used as one region – the range would then be about 10,000 km, like with the Americas.[5]

[5] In Asia, SNP haplotype heterozygosity is indicated to decrease whilst distance from Africa goes upward (Schlebusch et al., 2012).

**Text S3: Linearity**

Regarding heterozygosity, a non-linear pattern may be problematic when using linear regression to estimate where modern humans emerged (Liu et al., 2006) (assuming this origin is where they expanded from, Manica et al., 2007) – Liu et al. (2006) appear to have been treating the origin of the emergence and origin of the expansion as the same, so their observation regarding the potential problem also applies to estimations of where the expansion commenced. Indeed, the origin of the expansion may not be with the most diverse population, but geographically near to the most diverse (DeGiorgio et al., 2009). The Liu et al. (2006) observation would extend beyond linear regression to Pearson correlation, and to monotonic-based measures more generally.[6]

***Simulations: Increase in diversity over shorter distances?***

Previous research concerning the decline and linearity has featured simulations (e.g., DeGiorgio et al., 2009; Deshpande et al., 2009; Liu et al., 2006) – see Figure S8. Different sorts of simulations have been used (Deshpande et al., 2009). Deshpande et al. (2009) and DeGiorgio et al. (2009) distinguish between types of movement – i) *expansion* producing new populations, and ii) *migration* between adjacent populations which already exist due to expansion. Liu et al. (2006) combined these types (Deshpande et al., 2009). Deshpande et al. (2009) used both types, although not always. DeGiorgio et al. (2009) used a number of different types of simulations, including where both aforementioned types were used. In Liu et al (2006), a non-linear line was found regarding heterozygosity and distance. Indeed, Figure 4B of Liu et al. (2006) appears to suggest an increase in heterozygosity across shorter distances from the origin. A parallel pattern can be found in Figure 4A of DeGiorgio et al. (2009) (with, instead of geographical distance, the order in which populations are established). In simulations regarding that figure, both of the types of movement were used, with the migration rate being high (DeGiorgio et al., 2009). Indeed, whilst heterozygosity did decline, with a high (but not low) migration rate they found that heterozygosity is not at its utmost at the origin of the expansion, but reaches its highpoint nearby (DeGiorgio et al., 2009).

---

[6] Bearing in mind literature on correlation (Schober et al., 2018), in the context of expansion, non-linearity may be problematic outside of linear regression. Pearson's correlation has things alike with linear regression (Schober et al., 2018). The Pearson correlation coefficient measures a linear association (a type of monotonic association), whilst the Spearman correlation coefficient measures a monotonic association – a monotonic association is an association which is in one direction alone (e.g., on a graph, the variable on the *x*-axis increases with the *y*-axis variable) (Schober et al., 2018). Therefore, Figure 8C may have a non-monotonic quality. So, the possible issue is not non-linearity per se (because zero-order and semi-partial Spearman's correlations were also used in the present research in addition to Pearson's) but non-monotonicity. When using ranks (i.e., with Spearman), whether with heterozygosity being adjusted or not (Figure S5), it appears difficult to assess if heterozygosity rose across the shorter distances.

Liu et al. (2006) explain the non-linear pattern in their study as arising from greater migration into populations midway in the expansion. Through Deshpande et al. (2009), the Liu et al. explanation can be interpreted as referring to movement more generally (not just migration separate from expansion). DeGiorgio et al. (2009), regarding their own study, explain that diversity amongst incoming migrants is greater for populations midway than for populations at the origin and far-point of the expansion. Indeed, their simulations had been set-up where the population at the origin would only get migrants from later in the expansion, whilst the other populations (except the one at the far-point) can get migrants from earlier *and* further points along in the expansion (DeGiorgio et al., 2009). With more bottlenecks having been experienced midway than around the origin (so coalescence times may be longer around the origin), the utmost heterozygosity is somewhere between the origin and the midway point (DeGiorgio et al., 2009).

As described by Deshpande et al. (2009), in Liu et al. (2006) an expansion from the origin was along one type of route. DeGiorgio et al., (2009) also employed one type – see Figure S8A. Deshpande et al. (2009), however, represented two types of routes – one for expansion inside Africa (shorter route), and one for expansion to beyond Africa (longer route). These are represented, to some extent, in Figure S8B. The use of two types of routes was from the perspective of eastern Africa likely being where the expansion started (Deshpande et al., 2009). Figure 1 in Deshpande et al. (2009) presents results in which expansion and migration were used in simulation; whilst there was a linear fall in heterozygosity, the pattern did veer away from a linear pattern around the end of either type of route (Deshpande et al., 2009). Moreover, Figure 1 in Deshpande et al. (2009) would indicate that this particularly occurred for the shorter type of route (i.e., inside Africa). So, a pattern like in Liu et al. (2006) and DeGiorgio et al. (2009) likely would have been present if one type of route was used in Deshpande et al. (2009) like in those two other studies.

***Linear fall in diversity?***

On the relationship between distance from Africa and diversity, the relationship has been described as linear when linearity has been addressed (e.g., Prugnolle et al., 2005). Concerning the cranial form diversity of males, Betti et al. (2009) did find support for a non-linear relationship with distance from Africa (a model with linear and cubic terms, but not quadratic), although this was countered in the absence of two outlier datapoints. Regarding autosomal microsatellite heterozygosity, in Prugnolle et al. (2005), a linear fall in diversity was evident as distance from Africa increased, and gradients did not differ across groupings of populations (African, American, Asian, etc.) (Prugnolle et al., 2005). A non-linear relationship within Africa, if there is one, may have manifested as a difference in gradients. Yet, as noted in Text S2, Figures 1 and 2 in that study would indicate that not many populations were in any of the population groupings except the Asian one, so the ability to realise differences may not have been good.

A previous study using autosomal microsatellite heterozygosity (Balloux et al., 2009) may provide some suggestion of a non-linear relationship. Using HGDP-CEPH data, there was a decrease of autosomal microsatellite heterozygosity as distance from Africa grew in Balloux et al. (2009). Looking at Figure 3A of Balloux et al. (2009), perhaps there was actually an increase of heterozygosity at first (up to some distance under 5,000 km), and an overall quadratic pattern. This appears to be in line with Liu et al. (2006) Figure 4B and DeGiorgio et al. (2009) Figure 4A. However, going by populations, distances (Balloux et al., 2009), and Balloux et al. Figure 3A, only five populations were at distances less than 5,000 km. These HGDP-CEPH populations are in Africa (Tishkoff et al., 2009). Moreover, varying the location from which distances are measured leads to variation in the linear relationship between autosomal microsatellite heterozygosity and geographical distance (in terms of strength) (Ramachandran et al., 2005). Balloux et al. (2009), from the description in their methodology, did not necessarily select the peak point for autosomal microsatellite heterozygosity (for geographical distances to be from). So, a seemingly non-linear pattern may not necessarily reflect expansion, and instead happen due to the peak point not being selected.

***Monotonicity: Linear vs. quadratic***

Given that Figure 8C may provide some indication of a non-linear relationship, and also given the research reviewed above, the present research explored if there is a certain type of non-linear relationship between diversity and geographical distance, and, if so, re-estimated the origin of the expansion when using a non-linear model.[7] As for the type of model, it made sense to use a quadratic one (cf. Betti et al., 2009, regarding cranial diversity). This is because the pattern produced in DeGiorgio et al. (2009) (high migration) and line produced in the Liu et al. (2006) look like they are quadratic, and, because it appears that there may be a quadratic trend in Balloux et al. (2009) Figure 3A.[8]

The within-four-BIC benchmark (e.g., Manica et al., 2007) was employed to find whether any quadratic models predicted adjusted/unadjusted diversity more effectively than the best of the linear models. Quadratic models had the addition of a squared geographical distance term (the research of Betti et al., 2009, featured squared geographical distances). BICs can be used to estimate origin areas whether models are linear or non-linear (Betti et al., 2009); indeed, if the best quadratic model was

---

[7] Betti et al. (2009) estimated where the expansion originated both with the outliers (i.e., through non-linear models) and without them (linear models). Betti et al. (2009) used the Bayesian information criterion to select which model fits best (backward selection, including terms for climate); analysis in the present research was simpler, although with origin areas also indeed being estimated through non-linear models.

[8] Moreover, use of a quadratic term seemed sensible because the pattern in Betti et al. (2009) Figure 1A (retaining outliers) regarding cranial shape diversity at distances from Africa looks like it may be quadratic, despite inclusion of a quadratic term not being supported in their study.

more effective, then the benchmark was also to be used for estimating an origin area amongst quadratic models.

Two of the 106 sub-Saharan African populations had low diversity for their distance from Angola; these populations are Hadza and Mbororo Fulani (as they are called in Tishkoff et al., 2009) – when not adjusting heterozygosity, *z*-score-standardised residuals were found to be -5.31 and -3.41 respectively for Hadza and Mbororo Fulani, and, when adjusting heterozygosity, the standardised residuals were -5.34 for Hadza and -3.44 for Mbororo Fulani. Amongst linear models, analysis without these populations was not found to notably affect results (e.g., peak points in parametric analysis were unchanged). So, results for linear models were presented retaining these populations (except with respect to Figures 9 and 10). Whether the diversity of a population appears to be atypical could change depending on which location geographical distance is measured from. So, to define atypicality in general, one could ignore geographical distance. *Z*-scores not within the range of -3.29 to 3.29 can be regarded as outliers (e.g., Weng et al., 2022). Using that criterion (after transforming adjusted/unadjusted heterozygosity to *z*-scores), one can see that unadjusted heterozygosity is indeed low for the Hadza ($z$ = -4.77) and Mbororo Fulani ($z$ = -3.57) populations amongst the sub-Saharan African populations, and this is also apparent with adjusted heterozygosity for both Hadza ($z$ = -4.76) and Mbororo Fulani ($z$ = -3.60). Quadratic models were a better fit than linear models when using the 104 populations, but not when 106 populations were used. Therefore, the atypical datapoints were not included in the final analysis (i.e., using 104 populations) with respect to monotonicity and model fit.

**Table S1**

*List of Figures*

| Type | Figures |
|---|---|
| Africa map: Origin area? | 1A, 2A, 2C, 2E, 2G, 3B, 4C, 5A, 5C, 8A, 8B, 9E, 10A, S3A, S5B, S5D, and S7A |
| Africa map: Peak point(s) | 6, 7, 8A, 8B, 9B, 9C, 9E, 10, S5B, S5D, and S7A |
| Graphs: Associations/BICs | 1B, 2B, 2D, 2F, 2H, 3C, 4A, 4B, 5B, 5D, S2, S3B, S4, and S7B |

*Note*. In the present research, all Africa maps used coordinates from Betti et al. (2013).

**Figure S1**

*Cranial Diversity in Form and Shape Falls as Distance from Africa Greatens*

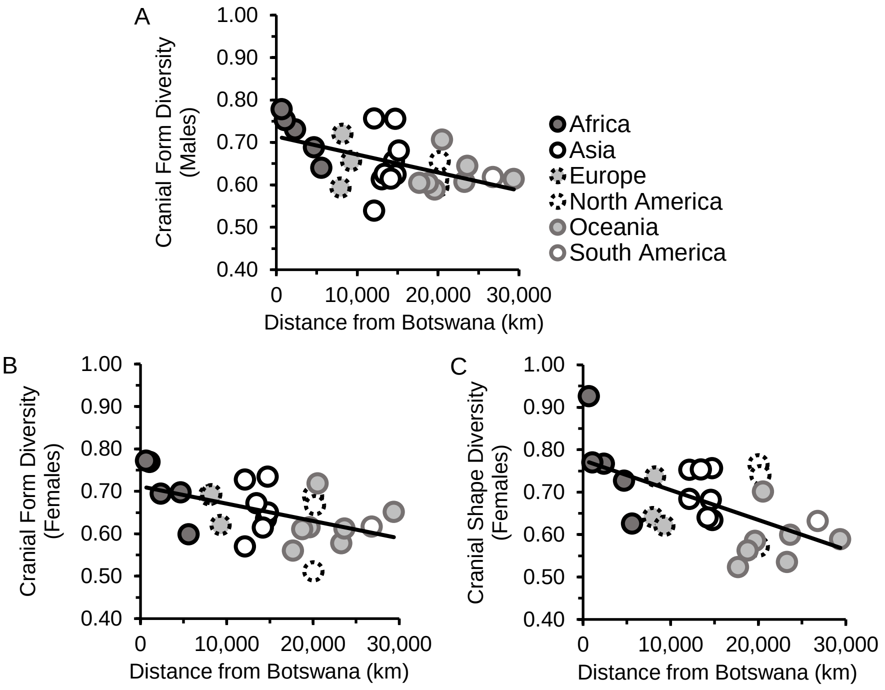

*Note*. Diversities are of the cranial form of males (S1A) in 28 populations, and of the cranial form (S1B) and shape (S1C) of females in 26 populations.

**Figure S2**

*Cranial Diversity and Geographical Distance*

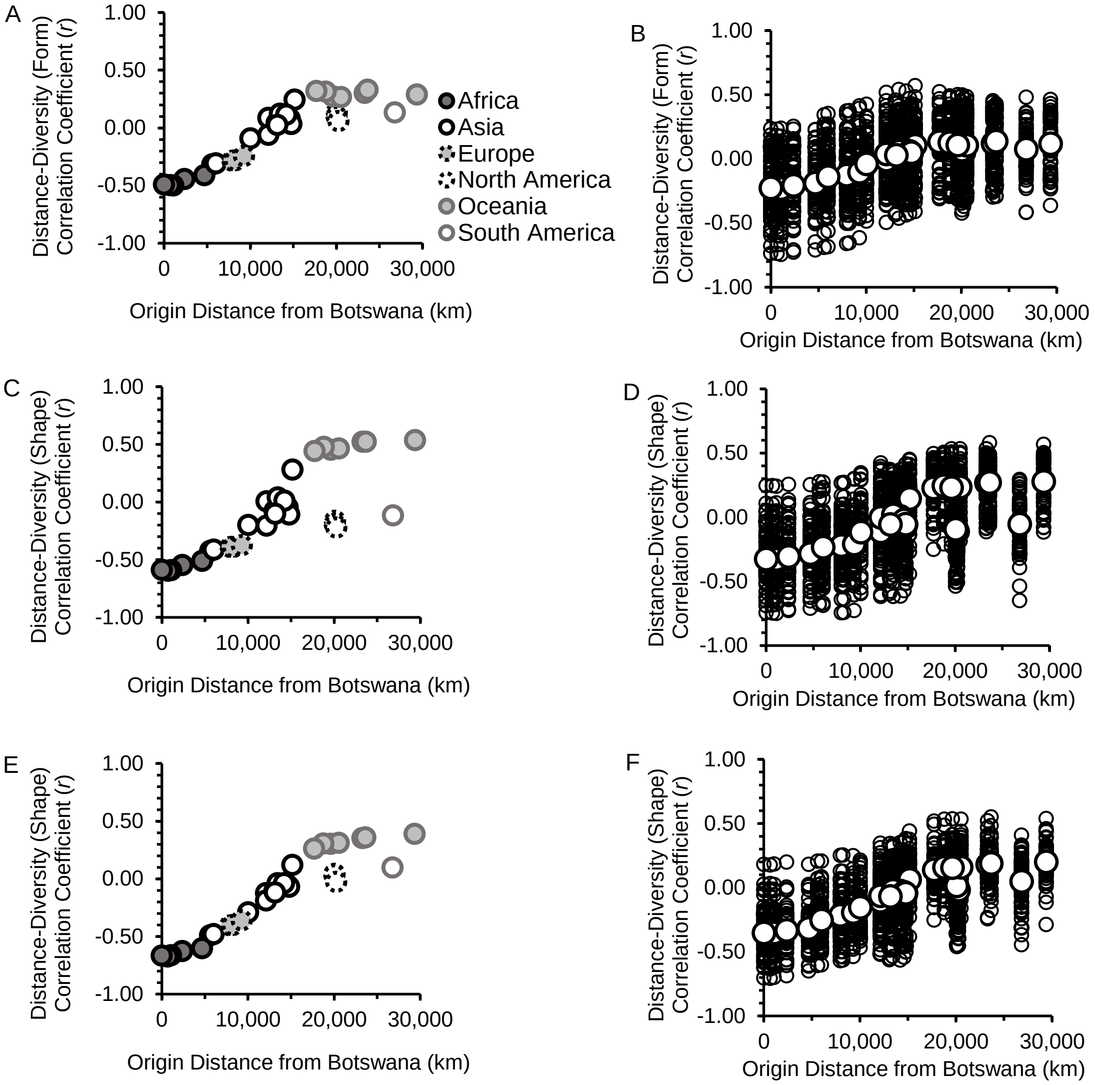

*Note*. Figures S2 and S4 show correlation coefficients presented on graphs like in Cenac (2022). These coefficients are for relationships between geographical distance and diversity within the cranium. Diversities are of the cranial form of females (S2A and S2B), the cranial shape of females (S2C and S2D), and the cranial shape of males (S2E and S2F). Panels on the left are from mean variances, and right panels are from variances inside individual (standardised) dimensions. In the right panels of Figure S2 (and in Figure S4), as in Cenac (2022), smaller circles are correlation coefficients, whereas bigger circles are the mean (average) of the coefficients concerning each origin. In order to reduce bias, coefficients can be transformed (Fisher *z*) and a mean (determined from those transformed values) itself transformed to an *r* correlation coefficient (Corey et al., 1998) – this process was used (as it was in Cenac, 2022) to arrive at the mean coefficients shown in Figure S2 right panels and the Figure S4 right panel.

**Figure S3**

*Cranial Size Dimorphism Adjusted for Absolute Latitude*

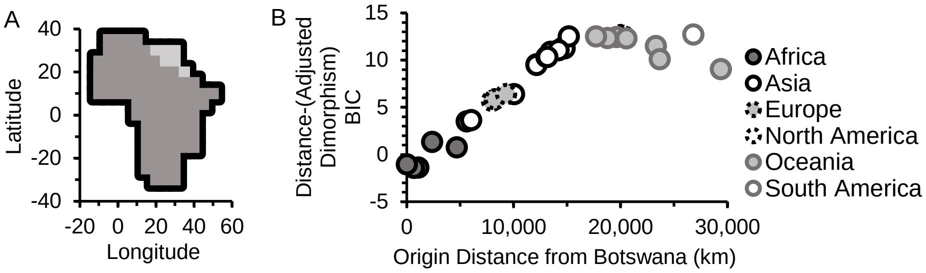

**Figure S4**

*Cranial Dimorphism*

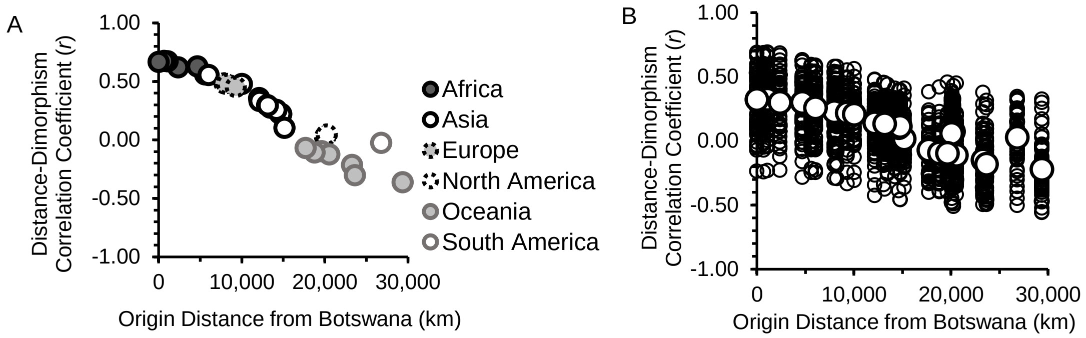

*Note*. In Figure S4A, the extent of relationships between geographical distance and cranial size dimorphism seems to have its highpoint when using an origin in Africa. As for Figure S4B, within cranial dimensions, the highpoint of correlation coefficients is also in Africa for relationships between distance and dimorphism. For Figure S4B, like in Figure S2 (right panels), the larger circles are means of coefficients achieved through the Fisher-z-transformation.

**Figure S5**

*Nonparametric: Autosomal Microsatellite Heterozygosity in Sub-Saharan Africa*

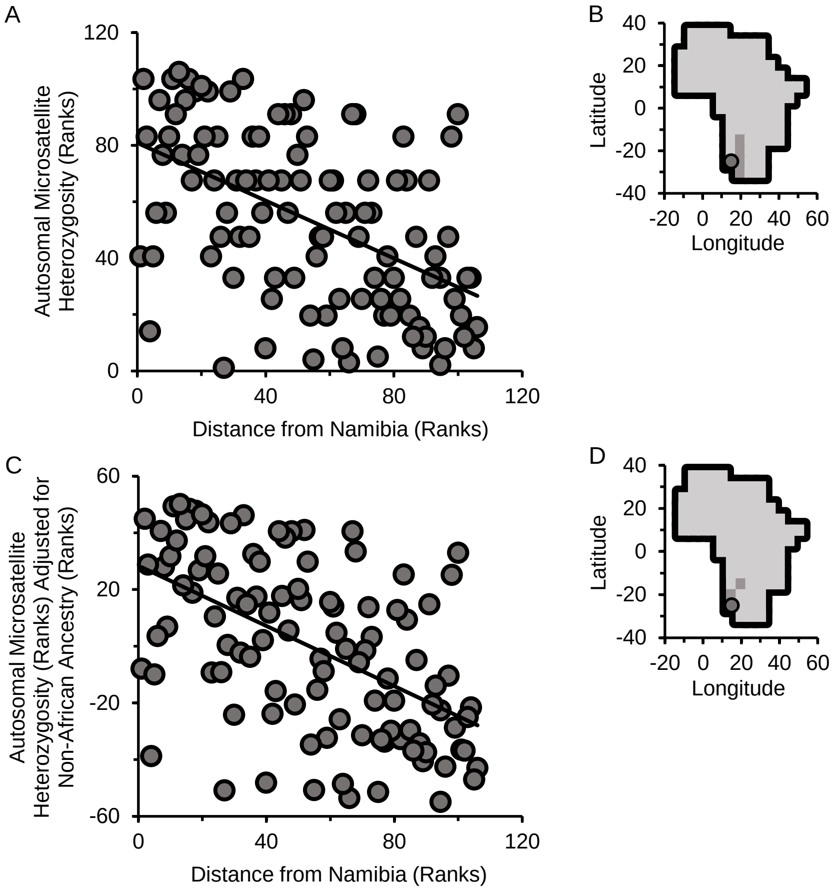

.

**Figure S6**

*Non-African Ancestry Within Sub-Saharan Africa*

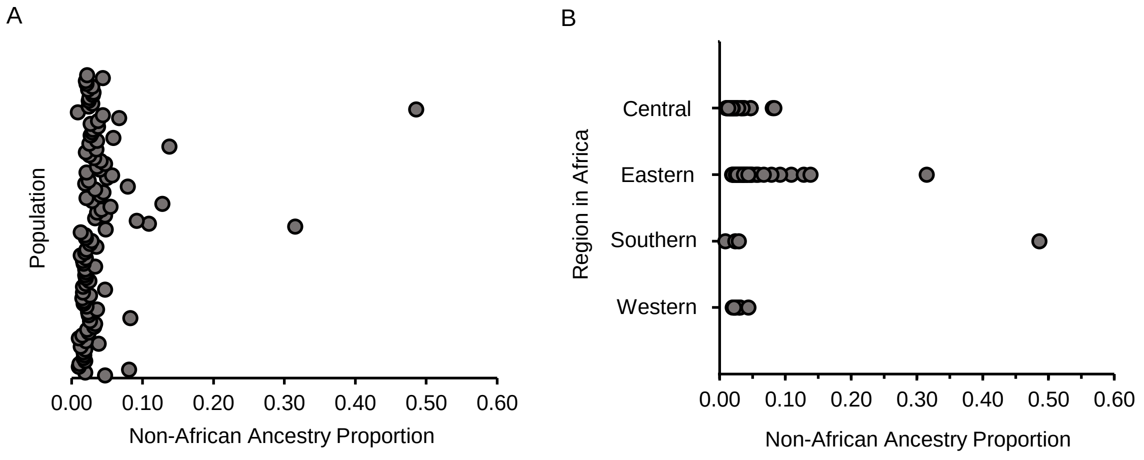

*Note*. Each datapoint is the non-African ancestry of a population in sub-Saharan Africa. Regions of populations were used from Tishkoff et al. (2009), and non-African ancestry was totalled from ancestry estimates in Tishkoff et al. (2009).

**Figure S7**

*Origin Area: Autosomal SNP Haplotype Heterozygosity in the Absence of San*

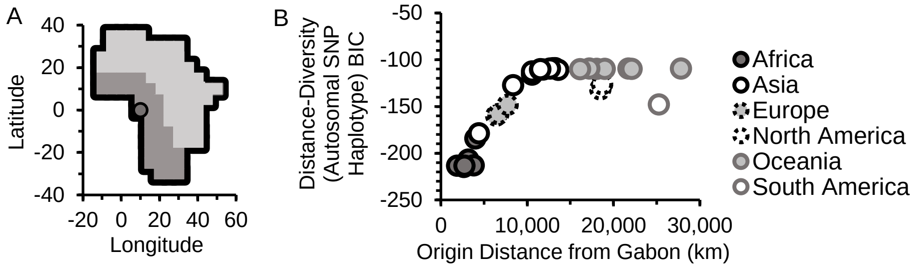

*Note*. Analysis with autosomal SNP haplotype diversity was done again without San – the peak point is at (0°N, 10°E).

**Figure S8**

*Types of Expansion Routes in Simulations (Regarding Previous Research)*

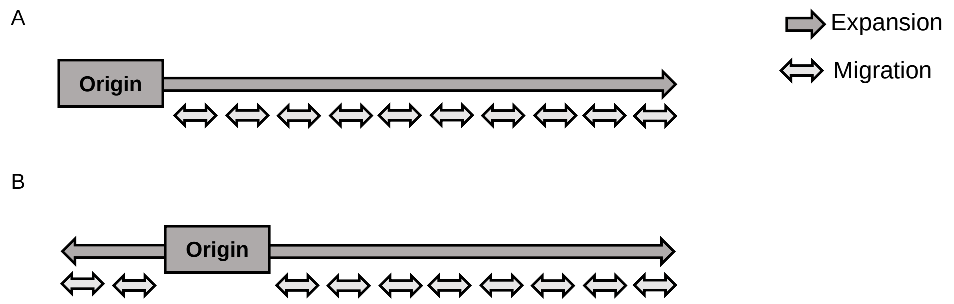

*Note*. Given previous research using simulations (DeGiorgio et al., 2009; Deshpande et al., 2009; Liu et al., 2006), Figure S8A is with respect to simulations in DeGiorgio et al. (2009) (simulations pertaining to their Figure 4A) and Liu et al. (2006), and Figure S8B is for Deshpande et al. (2009) (regarding their Figure 1). Expansion is distinguished from migration amongst next-door populations (Deshpande et al, 2009) in the figure (see Text S3 for more details). Representations are very general (e.g., the relative lengths of route types, number of populations) in Figure S8A and S8B. The present research (see *Results and discussion*) suggests that heterozygosity at shorter distances is more in line with Figure S8A than Figure S8B – i.e., in greater agreement with the Liu et al. (2006) and high-migration DeGiorgio et al. (2009) simulations.
